# AI mirrors experimental science to uncover a novel mechanism of gene transfer crucial to bacterial evolution

**DOI:** 10.1101/2025.02.19.639094

**Authors:** José R Penadés, Juraj Gottweis, Lingchen He, Jonasz B Patkowski, Alexander Shurick, Wei-Hung Weng, Tao Tu, Anil Palepu, Artiom Myaskovsky, Annalisa Pawlosky, Vivek Natarajan, Alan Karthikesalingam, Tiago R D Costa

## Abstract

AI models have been proposed for hypothesis generation, but testing their ability to drive high-impact research is challenging, since an AI-generated hypothesis can take decades to validate. Here, we challenge the ability of a recently developed LLM-based platform, AI co-scientist, to generate high-level hypotheses by posing a question that took years to resolve experimentally but remained unpublished: How could capsid-forming phage-inducible chromosomal islands (cf-PICIs) spread across bacterial species? Remarkably, AI co-scientist’s top-ranked hypothesis matched our experimentally confirmed mechanism: cf-PICIs hijack diverse phage tails to expand their host range. We critically assess its five highest-ranked hypotheses, showing that some opened new research avenues in our laboratories. We benchmark its performance against other LLMs and outline best practices for integrating AI into scientific discovery. Our findings suggest that AI can act not just as a tool but as a creative engine, accelerating discovery and reshaping how we generate and test scientific hypotheses.

## INTRODUCTION

Novel conversational artificial intelligence (AI) systems have tremendous potential to augment and accelerate biomedical discovery.^1,2,3,4^ However, it remains uncertain whether AI systems can propose creative, novel, and impactful hypotheses that rival those of senior scientists and meet the rigorous standards for publication in reputed scientific journals.

To explore this potential, we recently tested a novel AI system (named AI co-scientist) with a series of unsolved questions in biological and biomedical sciences. While the quality of the AI-generated hypotheses was impressive, verifying these hypotheses experimentally can require significant time and effort because they represent new areas of scientific inquiry requiring the planning and conduct of multiple new “wet lab” experiments. To more efficiently test the system, we therefore challenged its capabilities by using a specific unsolved question that had already intrigued our groups for over ten years and whose answer was recently uncovered through extensive experimental work, yet not disclosed publicly. At the time of testing the AI co-scientist, which occurred recently (Fig. 1), the experimental work answering the unsolved question used to challenge the AI system was confidentially under review in a scientific journal and no aspects of it were publicly accessible, thus ensuring that the AI system could not draw on prior knowledge of the correct answer when it was tested on this topic. This approach allowed us to directly assess the AI’s ability to generate plausible hypotheses by comparing its outputs to a newly known and hitherto-unpublished experimentally validated solution.

**Figure 1.**
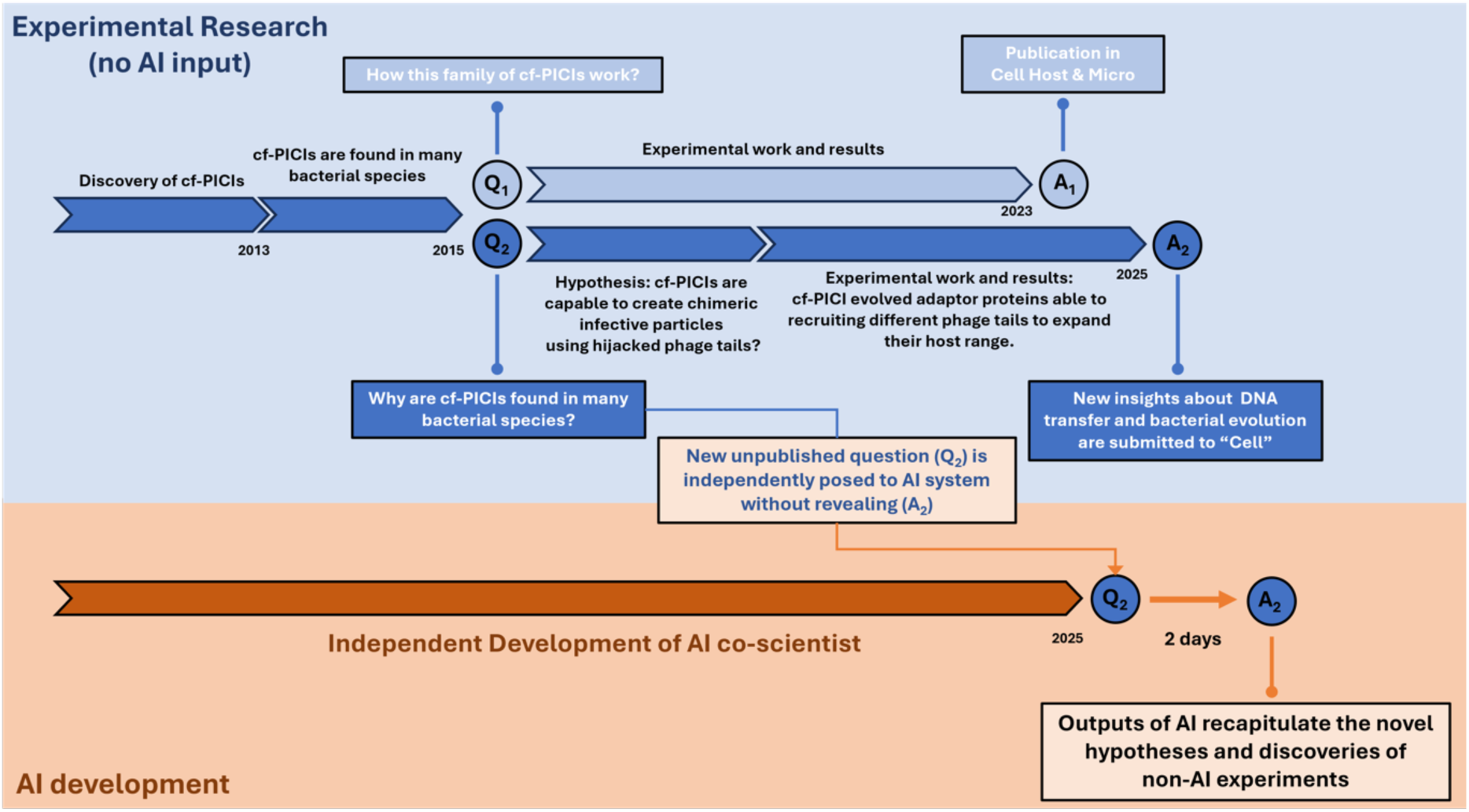
AI recapitulates the experimental discovery of a novel gene transfer mechanism. The blue section of the figure presents a flowchart outlining the timeline of the experimental research pipeline that led to the discovery of how cf-PICIs are mobilised between bacterial species. The orange section highlights the potential of AI to accelerate research by rapidly recapitulating, with no prior knowledge, the key experimental findings shown in blue.

Our work specifically focused on a unique family of phage satellites known as capsid-forming phage-inducible chromosomal islands (cf-PICIs).^5^ Phage satellites are mobile genetic elements that rely on helper phages for their lifecycle, including the formation of infective particles. Several families of phage satellites are well characterised,^6^ such as P4-like elements (in Gram-negative bacteria),^7^ phage-inducible chromosomal islands (PICIs; found in both Gram-positive and Gram-negative bacteria),^8,9^ PICI-like elements (PLEs; in *Vibrio cholerae*),^10^ and phage-inducible chromosomal minimalist islands (PICMIs; also in *Vibrio* species).^11^ These satellites depend entirely on helper phages to produce capsids and tails, which are critical for the packaging and dissemination of their genomes.

Remarkably, over a decade ago, we discovered a new family of satellites, which we named cf-PICIs (Fig. 1). Unlike classical PICIs and other satellites, cf-PICIs encode their own capsid-forming proteins, allowing them to produce small capsids independently of their helper phages.^5^ While these proteins are of phage origin, they have evolved to interact exclusively with cf-PICI components, exhibiting a specificity that prevents interaction with phage-encoded proteins and vice versa. This unique mechanism was published in 2023 after the concept of satellites as distinct genetic entities was well established (Fig. 1).^5^

Following their initial discovery, we observed an intriguing phenomenon: unlike other satellites, which are typically species-specific, identical cf-PICIs were frequently detected across different bacterial species. This observation was further validated through the development of methods to identify satellites in bacterial genomes.^12^ Given the narrow host range of phages and other satellites, we hypothesised that cf-PICIs employ an unprecedented mechanism of horizontal gene transfer to disseminate widely in nature. Over several years, we sought to uncover this mechanism and, through serendipity, recently pieced together the solution.

With this knowledge in hand, and confident that the scientific community was unaware of this breakthrough, we challenged AI co-scientist to hypothesise how identical cf-PICIs could be present in different bacterial species. We used this question as a test case to evaluate the system’s ability to answer scientific questions of interest. Since the system was not specifically trained in phage or satellite biology, or in bacterial evolution, we hypothesised that this would provide an unbiased test of AI’s ability to generate top hypotheses. The results were astonishing. AI co-scientist generated five ranked hypotheses, as reported in the result section, with the top-ranked hypothesis remarkably recapitulating the novel hypothesis and key experimental findings of our unpublished paper (Fig. 1, Table 1). The original manuscript, currently under review, is now available as a preprint to facilitate the understanding of the current work.^13^ This outcome demonstrates a fascinating and unsettling capability of AI systems to function at a level comparable to principal investigators, proposing hypotheses worthy of high-impact publication.

**Table 1.**
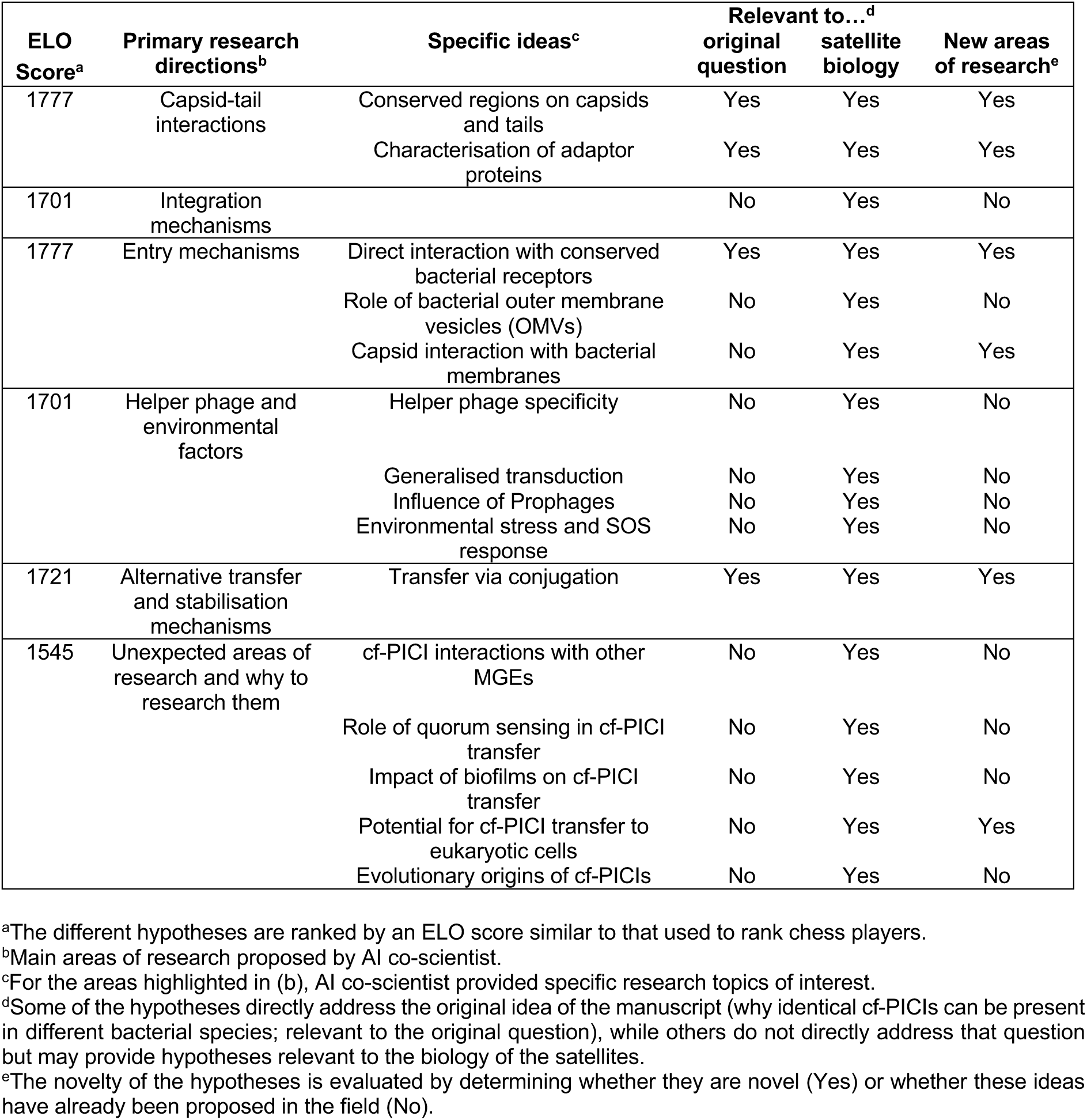
Summary of the AI’s provided hypothesis.

## METHODS

### AI System utilised for this challenge

As detailed elsewhere (https://storage.googleapis.com/coscientist_paper/ai_coscientist.pdf), the AI system utilised for this challenge comprised a LLM (large language model) driven novel hypothesis generation system, termed AI co-scientist. The AI co-scientist is a multi-agent system based on Gemini 2.0^14^ with inductive biases echoing the reasoning processes underpinning the scientific method. The system is designed to act as an AI collaborator for scientists and promotes an expert-in-the-loop workflow. Scientists can specify research goals in natural language with additional preferences and constraints reflecting their topic of expertise or experimental setup. They can also provide their own best-guess solution to the research or other feedback to guide the system. The system parses the research goal into a research plan configuration and employs a bunch of Gemini 2.0 based agents that continuously generate, debate and improve hypothesis and research proposals. These agents are specialised for generating, reviewing, ranking and improving the ideas with access to tools such as web search to enable literature review and summarisation. In particular, the system scales up test-time compute for scientific reasoning with an iterative, self-improvement loop, where the generated hypothesis competes in an Elo-score based tournament.^15^ The tournament accounts for multiple facets when scoring the proposed hypothesis and research directions including the criteria and constraints pre-specified in the initial research goal. The feedback and the win-loss patterns in the tournament are used to further refine and improve the ideas in the next iteration. At the end of the computation process, which can span for days instead of minutes with other frontier reasoning LLMs,^16,17,18^ the system comes up with a ranked list of hypotheses and research proposals that attempt to satisfy the provided research goal with an additional research summary overview.

In this specific challenge, only one “input document” was provided by expert scientists, as detailed in Supplementary Information 1, comprising only previously published and openly-available information as the research goal to the system. No specialist tools or proprietary databases were utilised by the system for this experiment. As introduced earlier, the proposed AI driven research directions are ranked by an ELO score similar to those used to rank chess players competing in tournaments.

### Framing the challenge as a prompt to the AI system: minimal input for maximum insight

We were extremely restrictive with the general information provided to the AI co-scientist system (Supplementary Information 1). Essentially, we supplied only a single page containing a brief background on phage satellites, including some references, highlighting two key papers. One manuscript described the original discovery of cf-PICIs,^5^ while the other detailed the development of a computational tool to identify phage satellites in bacterial genomes.^12^ Using rudimentary methodology, we had observed many years ago that identical cf-PICIs could be found in different bacterial species - a phenomenon we noted as the foundation of our unpublished experimental manuscript. The developed tool^12^ confirmed this observation and provided multiple additional examples of identical cf-PICIs present in different bacterial species.

To illustrate this observation, we included an example from the unpublished paper: two related cf-PICIs, PICIEc1 and PICIKp1, were identified in *Escherichia coli* GN02175, *Klebsiella pneumoniae* DSM30104, and other species, many of which are WHO priority pathogens. Specifically, PICIEc1 was detected in five genera and seven species, including *E. coli*, *K. pneumoniae*, *Shigella flexneri*, *Citrobacter freundii*, *Citrobacter amalonaticus*, *Enterobacter asburiae*, and *Enterobacter hormaechei*. Similarly, PICIKp1 was found in both *K. pneumoniae* and *E. coli*.

With this limited information, we posed the central question to the AI co-scientist system: why can cf-PICIs, but not other types of PICIs or satellites, be easily found across diverse bacterial species? What mechanism explains this phenomenon?

### Comparison of the AI co-scientist with other AI models

To enable a qualitative comparison with other AI systems, we also provided the same prompt as input to other state-of-the-art large language and reasoning models and a recently released “deep research” tool. This was done using their public facing user interfaces. The specific models compared were OpenAI o1 (https://openai.com/index/openai-o1-system-card/), Gemini 2.0 Pro Experimental (https://blog.google/feed/gemini-exp-1206/), Gemini 2.0-flash-thinking-exp-1219 (https://deepmind.google/technologies/gemini/flash-thinking/), OpenAI deep research (https://openai.com/index/introducing-deep-research/) and OpenAI o3-mini-high (https://openai.com/index/openai-o3-mini/).

## RESULTS

### AI-driven research directions for cf-PICI host range expansion

Based on the background information and existing hypotheses, AI co-scientist proposed five primary research directions to explain the broad host range of cf-PICIs, alongside a final point of potential areas of interest for future work. These were listed in order of preference and likelihood. A summary of the proposed hypotheses is presented in Table 1, and more detailed information about the generated output is provided in Supplementary Information 2. These correspond to:

1. **Capsid-tail interactions:** Investigate the interactions between cf-PICI capsids and a broad range of helper phage tails (ideas related to broad tail interacting, tail adaptor proteins, tail-binding sites, capsid-mediated interactions, etc).
2. **Integration mechanisms:** Examine the mechanisms by which cf-PICIs integrate into the genomes of diverse bacterial species (ideas related to integration, transposition, recombination, etc).
3. **Entry mechanisms:** Explore alternative cf-PICI entry mechanisms beyond traditional phage receptor recognition (ideas related to membrane vesicles, surface structures, membrane perturbation, etc).
4. **Helper phage and environmental factors:** Investigate the role of helper phages and broader ecological factors in cf-PICI transfer (ideas related to generalised transduction, prophages, community interactions, stress responses, etc).
5. **Alternative transfer and stabilisation mechanisms:** Explore other potential transfer mechanisms, such as conjugation, extracellular vesicles, and unique stabilization strategies, that might contribute to cf-PICI’s broad host range (ideas related to conjugative plasmids, EVs, membrane mimicry, unique stabilization strategies, etc).
6. **Unexpected areas of research and why to research them:** Beyond the core research directions outlined above, several unexpected areas might provide valuable insights into the broad host range of cf-PICIs

Below, we will evaluate the accuracy of the different proposed hypotheses. However, our initial review of the AI co-scientist outcomes left a strong impression - not only because one of the hypotheses (the most likely) accurately summarizes the findings of the unpublished paper,^13^ but also because the other hypotheses were plausible and, in some cases so compelling as to represent ideas that we plan to explore in the future.

### 1. Capsid-Tail Interactions

This hypothesis was ranked as the most promising, and to understand the magnitude of the outcome proposed by the AI system, it indeed summarised the experimental results presented in the accompanying paper.^13^ In that study, we demonstrated that cf-PICIs generate a novel biological entity consisting of cf-PICI DNA packaged into capsids formed by cf-PICI-encoded genes. These entities lack tails and are non-functional. We showed that they are produced even in the absence of helper phages, although their production significantly increases in their presence. Once cells carrying this new entity are lysed - through any mechanism disrupting cellular integrity - the tail-less cf-PICIs are released as inactive structures into the microbial community. There, they interact with tails from various phages infecting different bacterial species, forming functional and infective cf-PICI particles.

Since the host range of phages and satellites is determined by their tails, which specifically bind to structures on bacterial surfaces,^19^ this mechanism explains why identical cf-PICIs can be found in different species. Depending on the tail used to complete the cf-PICI infective particles, the cf-PICI DNA is delivered to different bacterial hosts.

Our experiments further demonstrated that some tail-less cf-PICI capsids could bind to multiple tails from phages infecting different species. This ability was linked to two proteins encoded by the cf-PICI genome: the tail adaptor and connector proteins. By swapping these proteins between two cf-PICIs, we showed that their expression determines which tails the cf-PICIs interact with and, consequently, which bacterial hosts their DNA can infect.^13^ Encouraged by these results, we have begun characterising the molecular mechanisms of these interactions using Cryo-EM.

The AI co-scientist’s suggestions for studying capsid-tail interactions were particularly relevant and insightful (see Table 1 and the paragraph entitled “What to Research in This Area” in Supplementary Information 2). Among the proposed ideas, all were plausible and non-falsifiable but two stood out and were extremely compelling:

a. *Conserved regions on capsids and tails*. Use Cryo-EM to identify conserved regions on cf-PICI capsids and phage tails that mediate their interactions. The AI system also recommended comparing these structures to pinpoint conserved contact points and using mutagenesis to test their role in binding and transfer.
b. *Characterisation of adaptor proteins*: Investigate the role of cf-PICI-encoded adaptor proteins in mediating interactions with diverse phage tails. It is also proposed to investigate the diversity of the genes encoding these proteins in the cf-PICI genomes.

It should be noted that our experimental paper also introduced novel concepts not identified by the AI system that were required to provide a complete picture of the new mechanism. For example, we reported for the first time that while helper phages are classically thought to induce satellite elements and provide all the components required for packaging, some helper phages induce the elements but fail to provide compatible tails, leading to the formation of tail-less cf-PICI capsids. Nevertheless, the manuscript’s primary finding - that cf-PICIs can interact with tails from different phages to expand their host range, a process mediated by cf-PICI-encoded adaptor and connector proteins - was accurately identified by AI co-scientist. We believe that having this information five years ago would have significantly accelerated our research by providing a plausible and easily testable idea.

### 2. Integration mechanisms

Even though this hypothesis does not directly address the original question, it must still be considered due to its importance once cf-PICIs are present in different species.

This hypothesis can be evaluated from two perspectives. On the one hand, while integration is undoubtedly essential for the stable maintenance of transferred mobile genetic elements in the recipient cell, it does not address the fundamental question of how identical cf-PICIs are found in multiple bacterial species. For integration to occur, cf-PICIs must first be mobilised across species, and the mechanism behind this inter-species transfer is not addressed here.

On the other hand, if we accept the first hypothesis as providing a plausible mechanism for cf-PICI delivery to different species, then the role of integration becomes highly relevant. One could envision a scenario where transfer occurs, but the transferred element fails to persist due to an inability to integrate into the recipient genome. As a case in point, a family of PICIs and cf-PICIs has recently been described that integrates into the late genes encoded by resident prophages.^20^ For these elements to integrate, the presence of the resident prophage in the cells is absolutely required. If the prophages cannot be mobilised to different species, that would impede the establishment (but probably not the transfer) of the cf-PICIs in other species.

Related to the cf-PICIs analysed in the related paper,^13^ our previous genomic analyses revealed that the attachment (*attB*) sites targeted by the cf-PICI integrases were highly conserved across diverse bacterial species. This conservation likely facilitates the integration of cf-PICIs following transfer. Notably, we did not include this information in our original challenge to the system, which makes this hypothesis even more intriguing, as it aligns with existing genomic evidence.

### 3. Entry mechanisms

This outcome was particularly intriguing. While these hypotheses were ranked with lower probability than the primary one, they explore novel pathways that could potentially explain how cf-PICIs are mobilised between species. It is important to emphasise that all these hypotheses operate under the assumption that the primary hypothesis - tail involvement in inter-species dissemination - was incorrect. In this section, the AI system strongly suggested confirming that tails were indeed not involved in cf-PICI dissemination (see Topic 4 in the section related to Entry Mechanisms, Supplementary Information 2). Based on this premise, the AI system proposed alternative mechanisms, some of which are genuinely exciting.

Importantly, although our experimental results indicated that these mechanisms are not the primary routes used by cf-PICIs for inter-species dissemination, we cannot entirely rule out the possibility that, on rare occasions, these events might occur. Below, we summarise the proposed ideas:

1. *Direct interaction with conserved bacterial receptors*. The AI’s first hypothesis, assuming tails were not involved, suggested testing whether tail-less cf-PICIs could directly interact with specific and conserved bacterial receptors. While there is no direct evidence supporting this idea, the presence of cf-PICI-encoded proteins with unknown functions made it worth considering. In fact, we investigated it in our experimental manuscript,^13^ and included strains that were either able or unable to produce tails compatible with tail-less cf-PICI capsids. These controls ultimately confirmed that tails are essential for transfer, effectively ruling out this mechanism.
2. *Role of bacterial outer membrane vesicles (OMVs)*. The second idea was equally compelling and has been proposed for the transfer of other satellites. Specifically, the Tycheposons, a satellite family found in *Prochlorococcus*,^21^ have an unknown transfer mechanism. It has been hypothesised that some of these elements are mobilised between microbial communities via vesicles, while others are transferred in viral particles.^21^ This dual mode of transfer might explain their remarkable genetic diversity and the unusual range of sizes for these satellites (4–200 kb). Although we found no direct evidence for this mechanism in cf-PICIs, the hypothesis remains an exciting avenue for future research.
3. *Capsid interaction with bacterial membranes*. The final hypothesis suggested by AI co-scientist proposed that cf-PICI capsids could directly interact with and penetrate bacterial membranes, bypassing receptor-mediated entry mechanisms. This might involve membrane fusion or pore formation for entry. To investigate this, the AI recommended experiments using liposome model systems to study interactions between purified cf-PICI capsids and artificial membranes with varying lipid compositions. Techniques such as cryo-EM and fluorescence microscopy were suggested to visualise these interactions and assess for evidence of membrane fusion or pore formation (see Topic 3, section Entry Mechanisms, Supplementary Information 2).

In summary, taken out of context, these ideas might seem implausible. However, in light of the possibility that the primary hypothesis could be incorrect, these alternative hypotheses are highly creative and, more importantly, straightforward to test. This is crucial. The AI system provides innovative ideas that have the potential to revolutionise our understanding of how satellites function. Furthermore, the initial testing of these ideas is relatively simple and grounded in a clear rationale. In our experience, the necessity of including controls to confirm the role of tails in cf-PICI transfer demonstrates the importance of these alternative mechanisms. Even though our findings did not support these hypotheses as primary routes, they remain valuable for exploring the broader context of satellite dissemination and inter-species gene transfer.

### 4. Helper phage and environmental factors

In this section, the AI co-scientist proposed several interesting ideas. However, these ideas were not entirely novel and, in some cases, have already been suggested for the transfer of other satellites. Moreover, many of these ideas are not specific to cf-PICIs and could be applied to other satellites. As a result, while they provide valuable insights into satellite transfer mechanisms, they do not directly address the unique question of why identical cf-PICIs, but not other satellites, are frequently found in different bacterial species. Nevertheless, the hypotheses are thought-provoking and offer avenues for further exploration.

1. *Helper phage specificity*. The first hypothesis suggested investigating the range of helper phages capable of supporting cf-PICI transfer and whether helper phages with broad host ranges could explain the interspecies dissemination of these elements (Table 1, Supplementary Information 2). Interestingly, this idea has already been validated for classical PICIs in *Staphylococcus aureus*. Research, including our own, has shown that helper phages for classical PICIs can inject their DNA into a range of *Staphylococcus* species, such as *S. epidermidis*, *S. xylosus*, and *S. chromogenes*, and even into unrelated species like *Listeria monocytogenes*.^22,23,24^ This broad host range could explain the presence of SaPI-like elements in different species. However, contrary to cf-PICIs, which are found as identical elements in multiple species, identical SaPI elements have not been observed across species. This suggests that while the mechanism may facilitate transfer, as occurs in the laboratory, these elements are not maintained in recipient cells, likely because they do not confer additional benefits. Integration stability and other factors may also play a role here, as previously hypothesised.
2. *Generalised transduction*. The second idea relates to generalised transduction, where phages inadvertently package host DNA instead of their own.^25^ This could theoretically enable cf-PICI transfer without requiring specific interactions between cf-PICI capsids and phage tails. This mechanism has already been demonstrated for *S. aureus* PICI SaPIbov2.^22^ However, as with helper phage specificity, this hypothesis does not explain why cf-PICIs are uniquely found in multiple species while other satellites are not. Since cf-PICIs and classical PICIs share similar sizes and genomic loci, it remains unclear why other satellites would not also benefit from these mechanisms.
3. *Influence of prophages*. Another hypothesis suggests that prophages may play an important role after cf-PICI transfer by providing *attB* sites for integration or by acting as helper phages in the new species (Topic 3, Supplementary Information 2). While this idea is plausible and aligns with existing knowledge,^20^ it does not address the critical question of how cf-PICIs initially transfer between species. Once transfer occurs, it is reasonable to assume that cf-PICIs might interact with new helper phages to promote further dissemination in the recipient species, and these interactions must be investigated, as we did in our experimental paper. However, the mechanism enabling the initial transfer remains unexplained.
4. *Environmental stress and SOS response*. The final idea, outlined in Topic 4 (Supplementary Information 2), proposed that environmental stressors, including those triggering the SOS response, could induce resident prophages and thereby facilitate satellite transfer. While this concept is broadly applicable to satellite mobilisation, it does not specifically address the unique dissemination of cf-PICIs. This hypothesis is more general and represents a common mechanism of phage and satellite activation under stress conditions.

In summary, as these hypotheses become broader and less specific, they lose focus on the primary question that initiated this study: why identical cf-PICIs, but not other satellites, are found across different bacterial species. Despite this limitation, these general ideas remain valuable and merit consideration. They align with previously published concepts and provide a broader framework for understanding satellite transfer. Although they do not fully resolve the unique case of cf-PICI dissemination, they offer a foundation for further exploration and may guide future studies into the complex interplay between satellites, phages, and their environments.

### 5. Alternative transfer and stabilisation mechanisms

One of the significant advantages of using AI systems is their ability to propose research avenues in ways that differ from human scientists. A compelling example of this is the first hypothesis presented in this section. Topic 1 suggests exploring conjugation as a potential mechanism for satellite transfer. This idea is particularly exciting and has never been considered by investigators in the field of satellites.

In a manner analogous to how some satellites integrate into prophages, it is plausible that satellites could integrate into conjugative elements such as conjugative plasmids, integrative-conjugative elements (ICEs), or integrative mobilisable elements (IMEs). This integration would enable satellites to be mobilised via conjugation. Given that conjugation is generally more promiscuous than phage-mediated transfer, this hypothesis is currently under investigation in our lab.

Further related to conjugation, the AI system proposed that cf-PICIs (and potentially other satellites) could hitchhike on conjugative elements for transfer. In addition to integrating into these elements, satellites could potentially carry an *oriT* sequence in their genomes. These *oriT* sequences are recognised by the conjugative machinery to initiate the transfer process.^26^ If satellites encoded such sequences, they could hijack the conjugative machinery to facilitate their mobilisation between species.

Supporting this idea, it is worth noting that some SaPIs carry *cos* sequences, which enable them to hijack the packaging machinery of *cos* phages.^23,27,28^ Much like *oriT* in conjugation, *cos* sequences are recognised by the phage machinery to initiate packaging, ensuring efficient transfer. The possibility that satellites could similarly exploit *oriT* sequences to hijack conjugation is an exciting and innovative hypothesis. This idea, directly inspired by the AI system and not previously proposed by human researchers, is now being actively explored in our lab. In collaboration with Prof. Eduardo Richa’s group at the Pasteur Institute, we have identified potential satellite candidates with *oriT* sequences and are currently investigating their role.

The remaining hypotheses in this section were less groundbreaking. Topic 2 revisited extracellular vesicles as a mechanism for satellite transfer, a concept previously proposed here and in other contexts. Topic 3, focusing on stabilisation mechanisms, explored the role of the immune system in facilitating satellite entry into new cells but did not provide a strong link to inter-species transfer. Topic 4, which discussed alternative replication strategies, was less compelling. cf-PICIs utilise a replication strategy identical to classical PICIs,^5,8^ making this idea less relevant to the question at hand. Additionally, while stabilisation (Topic 3) and replication strategies (Topic 4) might play a role after transfer, their relevance to the initial inter-species dissemination of cf-PICIs remains unclear.

In conclusion, the first hypothesis - conjugation as a potential mechanism for cf-PICI transfer - stands out as innovative and exciting. By contrast, the other ideas presented in this section were less relevant and often repetitive or diffuse. The strength of the AI system lies in its ability to generate fresh and unbiased hypotheses, and in this case, it delivered a novel and promising avenue that we are actively pursuing. This hypothesis underscores the value of AI-generated ideas, not only for advancing our understanding of cf-PICI transfer but also for proposing additional principles that may impact satellite biology.

### 6. Unexpected areas of research and why to research them

The topics proposed in this section are undoubtedly of interest and align well with satellite biology, including cf-PICIs. However, they do not directly address the original question posed in this study. It appears that the AI system intended to expand the scope of research by highlighting broader areas beyond the specific focus of our inquiry.

The first proposed area relates to *cf-PICI interactions with other MGEs*. This is a highly relevant and active area of research in satellite biology. Both we and others have been investigating the impact of satellites on their helper and non-helper phages,^29–32^ as well as other MGEs such as plasmids and other satellites. For instance, we demonstrated that some PICIs use other PICIs for induction.^33^ Moreover, ww have also shown that PICIs and phages influence the size of non-conjugative plasmids that rely on transduction for transfer.^34^ This line of inquiry holds enormous potential, and some members of our team are already exploring it further.

The second topic suggests examining the *role of quorum sensing in cf-PICI transfer*. Recent studies have revealed an important role for quorum sensing in phage biology.^35^ Extending this concept to satellites is logical, given that satellites rely on phages for induction and transfer. This hypothesis could open new avenues for understanding the regulatory mechanisms of satellite mobilisation.

The third idea proposes analysing the *impact of biofilms on cf-PICI transfer*. This area remains largely unexplored but has significant potential. Biofilms are known to enhance gene transfer^36^ and often consist of diverse bacterial species, providing a plausible niche for inter-species satellite transfer. Remarkably, SaPIbov2 encodes Bap,^37^ a protein involved in biofilm formation,^38^ further supporting this idea’s relevance.

The fourth suggestion, examining the *potential for cf-PICI transfer to eukaryotic cells*, seems less biologically relevant. To date, no satellite sequences have been identified in eukaryotic cells, making this hypothesis speculative and less impactful for the current study.

The final topic addresses the *evolutionary origins of cf-PICIs*, a classic question often raised in discussions about satellites. While this is undoubtedly an important and exciting area of research, it diverges from the original scope of this work.

The AI-generated ideas in this section highlight exciting and diverse research areas that could advance our understanding of satellite biology. While they do not directly address the primary question of cf-PICI inter-species transfer, they underscore the broader significance of satellites and their interactions with other biological systems.

#### Analysis of other LLM systems

Having been impressed by the outcomes generated by the AI co-scientist system, we were intrigued to see what kind of hypotheses would be provided by other recently available Large language and reasoning models as well as “deep research” agentic systems, which were challenged using the same document that was used to challenge the AI co-scientist (Supplementary Information 1). These include OpenAI o1 (https://openai.com/index/openai-o1-system-card/), Gemini 2.0 Pro Experimental (https://blog.google/feed/gemini-exp-1206/), Gemini 2.0-flash-thinking-exp-1219 (https://deepmind.google/technologies/gemini/flash-thinking/), OpenAI deep research (https://openai.com/index/introducing-deep-research/), and OpenAI o3-mini-high (https://openai.com/index/openai-o3-mini/). Since a very specific question was posed in these analyses, we did not perform this competition to rank the different models but rather to observe the type of thinking they developed. It is worth noting that these systems were reasoning and using compute that spanned in the order of seconds and minutes and we took only one sample from the system. The co-scientist on the other hand, performs iterative self-improvement and reasoning that can span days to come up with its solution. Additionally, the AI co-scientist does use Gemini 2.0 models as specialised agents in the system. The AI co-scientist is a flexible system and can easily incorporate the other AI systems compared here as agents or tools, so they are not necessarily competing with each other. However, we thought it was still helpful to highlight the differences.

The results, summarised in Table 2, indicate that, contrary to the AI co-scientist system, none of the other AI systems were able to recapitulate the results of the experimental manuscript. However, they proposed some ideas, many of which were also suggested by the AI co-scientist. Supplementary Information 3 contains the outputs generated by the different AI systems, which are summarised below.

**Table 2.**
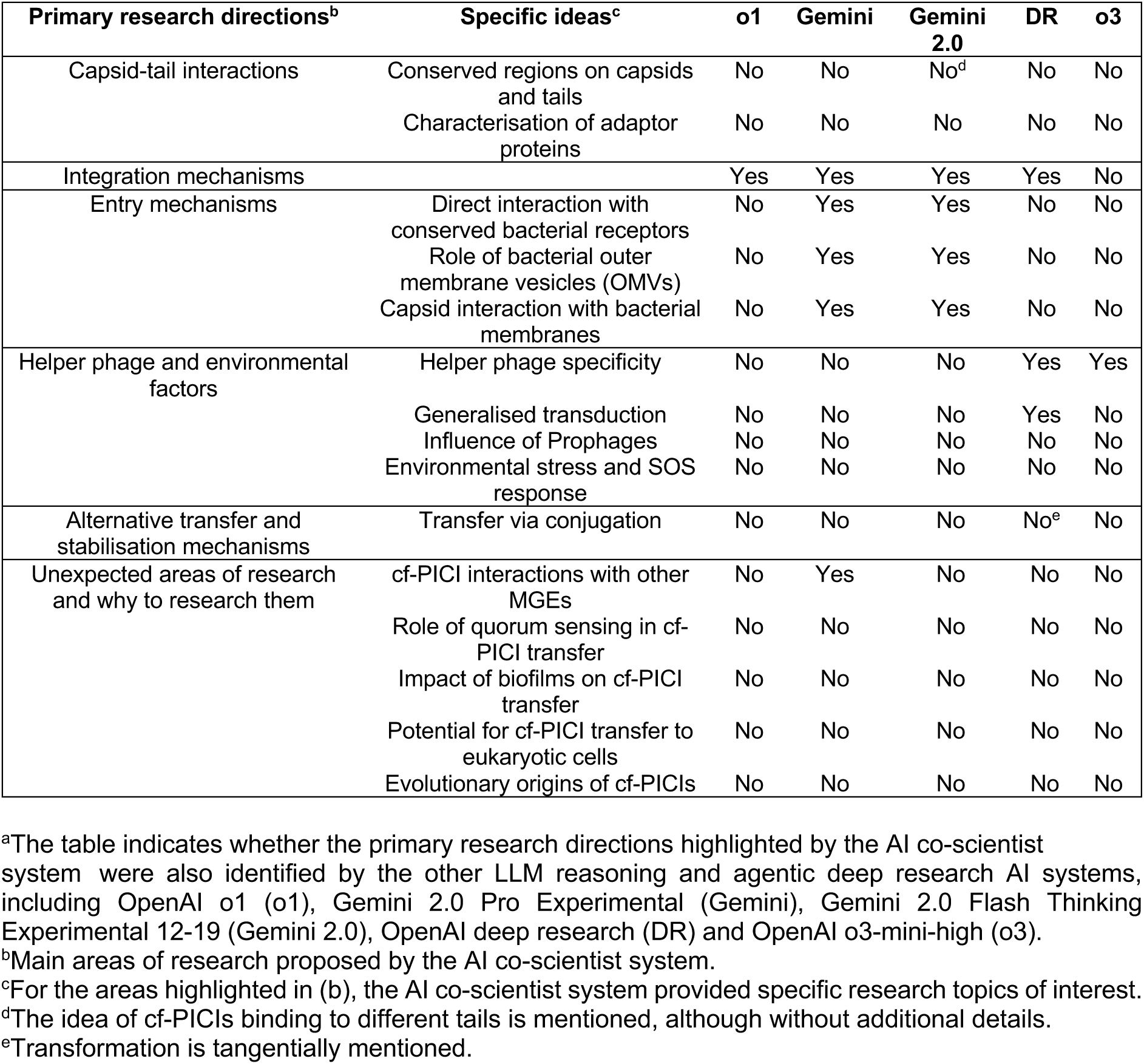
Comparison of the different AI systems^a^.

#### OpenAI o1

The first idea provided by this system, entitled *’The Minimal Dependence on Helper Phage Tails Enables Broad “Helper”*,’ seemed to identify the mechanism explaining the broader distribution of cf-PICIs between different species. However, the wording was a bit confusing, and after carefully analysing it, the hypothesis proved to be incorrect. OpenAI o1 hypothesised that tail genes are more conserved than capsid genes, and therefore, the same tails can be used for many capsids, even for unrelated capsids. In fact, as summarised at the end of the report, OpenA o1 hypotheses that ‘because cf-PICIs provide their own head morphogenesis and genome-packaging modules yet borrow a tail module widely conserved among many phage “helpers,” they can efficiently transfer between taxonomically distant bacteria. This unique autonomy of capsid and packaging - combined with broad phage-tail compatibility - drives their cross-species spread.’

However, even though the idea of combining tails and capsids may seem correct, it is not in this case. Tails define the tropism, and if several phages use the same tails, they will have the same tropism, so their DNAs will be present in the same species. Tail genes are not more conserved than capsid genes, and each phage has a combination of these, which determines the compatibility of their capsids and tails. The correct answer is not the use of the same tail with different elements, but rather the opposite: the use of different tails, with different tropisms, by the same cf-PICI.

OpenAI o1 also mentioned mechanisms related to integration, which, as previously discussed, are necessary for transfer once the elements have arrived in the new species. It also suggested that the fact that cf-PICIs package their DNA independently of their helper phages may be important to overcome the species barrier, but the rationale behind this seems incorrect. Note that some satellites (cf-PICIs) encode their own packaging mechanisms,^13^ which allow them to preferentially package their own DNA instead of the helper phage DNA.

#### Gemini 2.0 Pro Experimental

This model proposed two main ideas that were also suggested by the AI co-scientist system. The first relates to the potential ability of tail-less cf-PICIs to interact, via different mechanisms, with bacterial receptors in a way that does not require tails. The second idea suggested exploring mechanisms of cf-PICI integration once the transfer had occurred. As previously mentioned, so far it seems clear that tails are required for cf-PICI transfer, while the mechanisms involved in the integration of the elements in the recipient cells, though not directly addressing the main question, are definitely relevant to the biology of the phage satellites.

#### Gemini 2.0 Flash Thinking Experimental 12-19

As with Gemini 2.0 Pro Experimental, the main hypothesis suggested by this Gemini Thinking model related to the idea that tail-less cf-PICIs can directly interact with different bacterial structures present in different species, explaining why these elements can be mobilised between species (hypothesis 1: Relaxed Host Specificity of cf-PICI Capsid Receptor Binding). Compared to the AI co-scientist, and in addition to outer membrane proteins, this system even included additional potential receptors, such as the lipopolysaccharide (LPS) core regions or peptidoglycan components.

This Gemini Thinking model also mentioned integration as a key factor to be analysed in answering the question of inter-species cf-PICI transfer (hypothesis 3: Post-Entry Survival and Integration in Diverse Hosts). Related to this, the system also proposed ideas not previously suggested, such as analysing the impact that the immune systems present in the recipient species may have on controlling cf-PICI transfer.

Gemini 2.0-flash-thinking exp-1219 also proposed an additional line of investigation (hypothesis 2: “Universal” Packaging and Tail Compatibility), which included two main ideas. One relates to the fact that cf-PICIs may be less dependent on host factors to package their DNA into their assembled capsids. There is no evidence of this. Importantly, in this aim, it was also suggested that “the interface between the cf-PICI capsid and the helper phage tail might be less stringent than in typical phage systems. This could allow cf-PICIs to utilie tails from a broader range of helper phages, even those from distantly related bacteria. This could be due to a simpler, more generic interaction mechanism at the capsid-tail junction.” This definitely fits with the mechanism that explains how cf-PICIs spread inter-species. However, this result was less spectacular than that reported by the AI co-scientist because, overall, the report proposed that the mechanism of inter-species transfer depends on the capsid, not the tail. In fact, in the section *Novelty and Specificity of the Mechanism*, it is indicated that “This mechanism is novel because it highlights the capsid itself as the primary driver of broad host range for cf-PICIs, contrasting with the typical view where phage host range is primarily determined by tail fibre specificity and host factors.” This suggests that the system assumes inter-species transfer occurs via capsid interaction and probably, intra-species transfer occurs via interaction with different tails. In any case, it would have been nice to know how we would have interpreted this in a scenario where we did not know the correct answer. It is tempting to speculate that even indirectly, maybe these ideas would have served to initiate a discussion that would eventually lead to the identification of the right answer.

#### OpenAI deep research

This system provided an extensive overview of the system, including general concepts relevant to the biology of the cf-PICIs and other satellites. The complete report can be found in Supplementary Information 4. The system also provided a very clear hypothesis about how cf-PICIs move between species. This detailed pathway has been incorporated into Supplementary Information 3.

In this report, the system proposed that the inter-species transfer of cf-PICIs depends on phages that can infect multiple bacterial species. The report states that “cf-PICIs across species likely exploit phages that serve as ‘bridges’ between different bacteria. Indeed, broad-host-range or generalised transducing phages could carry a cf-PICI from one species to another in a single transduction event. Even phages with a narrower range might facilitate stepwise transfer – for instance, a PICI could move from species A to closely related species B via one phage, and from species B to species C via another phage that infects B and C, and so on, eventually appearing in distantly related hosts.”

This idea was also proposed by the AI co-scientist, and as mentioned before, it works in the laboratory with the classical PICIs from *Staphylococcus aureus* (SaPIs) but does not seem to be relevant in vivo, since SaPIs are exclusive to *S. aureus*, with identical elements not being present in other species. OpenAI deep research does not recapitulate the main mechanism that explains inter-species cf-PICI transfer, although it does emphasise the relevance of the integration mechanism in the success of the transfer. Tangentially, the system also mentions the unlikely role that other mechanisms of gene transfer, such as transformation, may play in this process. There is no evidence supporting this idea.

#### OpenAI o3-mini-high

Contrary to other AI systems, OpenAI o3-mini-high provided only a hypothesis with a promising title: “*Autonomous Capsid Assembly Coupled with Promiscuous Tail Exploitation*” (Supplementary Information 3). However, upon closer examination, the system proposes that “many temperate phages have tail proteins that are more conserved and functionally promiscuous across diverse bacterial species.” This idea was also suggested by other systems and, as previously mentioned, is incorrect. The issue is not that the same tail can be used to inject cf-DNA into different species, but rather that the tail-less packaged cf-PICI DNA can interact with different tails, each one specifically infecting a single species.

## DISCUSSION

AI has illustrated groundbreaking potential for dramatic improvement in biological research at points along the experimental pipeline and has numerous opportunities to reshape the end-to-end scientific process. With regards to hypothesis generation, our investigation into cf-PICI dissemination demonstrated that generated co-scientist-assisted hypotheses were found to be creative and logically reasonable. The end result was a range of novel and exciting approaches that were experimentally testable. The system’s ability to generate customisable hypotheses quickly and efficiently is particularly striking, as it provides a gateway to enhance the speed of scientific discovery. The ideas embodied the high standards of cutting-edge research while demonstrating the advantages of pushing beyond traditional paradigms, suggesting alternative mechanisms for cf-PICI transfer which opened up new research avenues. The co-scientist system has the ability to search and disseminate a much wider research literature space, presenting the opportunity to build transdisciplinary bridges across numerous research disciplines in a way that has never before been possible. The system’s suggestions, such as the discovery of how cf-PICIs are transferred inter-species or the conjugation hypothesis for satellite transfer, represent genuinely innovative avenues of research that had not been considered in the field of satellite biology. Given the speed with which these ideas were generated and their logical coherence, it is clear that this system could be a tool used by researchers in the biological sciences.

Here we not only focused on the AI co-scientist system but also compared different state-of-the-art LLM reasoning and agentic deep research AI systems, and they generated a range of distinct outcomes. Only one system (AI co-scientist) recapitulated in detail the findings of our experimental study, demonstrating an ability to converge on established mechanisms. Several introduced new avenues of research, suggesting hypotheses that had not been previously considered. Additionally, some ideas emerged across multiple systems, yet they did not directly answer the core question of our study. This raises an intriguing consideration: while one could combine insights from multiple systems to explore a broader spectrum of possibilities, the recurrence of a particular idea across different models does not necessarily indicate that it is the most accurate or scientifically robust explanation. Our analysis indicates that AI systems vary in their capacity for creativity, hypothesis generation, and problem-solving approaches. Some may be more conservative, favouring established patterns, while others may introduce more speculative but potentially groundbreaking concepts. Therefore, consistency among AI-generated hypotheses should not be equated with validity or superiority; rather, it highlights the need for careful evaluation and experimental validation to discern which ideas hold true biological relevance.

The AI co-scientist is not only a tool for hypothesis generation but also a catalyst for accelerating research progress from numerous angles. For example, the co-scientist can be utilised to ascertain combinatorial questions, such as questions surrounding drug repurposing or the generation of animal disease models. The co-scientist can also perform backward-looking queries, such as determining plausible explanatory hypotheses based on a given dataset. The ability to conduct both backward- and forward-looking analyses is extremely vital in the current age of high-resolution, high-throughput data generation technologies, such as next-generation sequencing, single-cell transcriptomics, and novel protein sequencing methodologies, since it will allow researchers to evaluate gathered datasets in ways that have not yet been previously considered. Importantly, while AI-driven approaches can generate valuable hypotheses and insights, experimental validation by scientists remains essential to confirm and refine these predictions, ensuring their biological relevance and real-world applicability.

However, with every rose, there is a thorn, and there are some important challenges associated with the integration of AI in scientific research. While numerous plausible solutions can rapidly be generated, researchers must now move their thinking towards efficient and cost-effective frameworks to evaluate many putative approaches. Intellectual property could also present itself as a challenge. With AI-assisted hypothesis generation, scientific thinking may become more democratised, and the process of idea attribution to specific individuals or research teams may need to be reconsidered. Additionally, the muscle of creative and critical thinking among scientists must be preserved. AI systems present the opportunity for over-reliance, which may result in researchers pursuing avenues that are not as relevant or impactful as they initially appear. While the AI system can propose creative ideas, it lacks the nuanced understanding that comes from years of expertise and experience in the field. This could lead to the pursuit of hypotheses that, although interesting, might not yield significant results or contribute meaningfully to the research question at hand.

The critical impact of the ability to rapidly produce plausible, mechanistic hypotheses that are supported by experimental findings can easily be scaled and tested to other critical questions in biomedical research. This paradigm-shifting innovation opens numerous questions, such as how to rapidly evaluate numerous hypotheses, the nature of intellectual property rights, evolutions to the grant application process, and the challenge of distinguishing independent contributions in collaborative environments involving AI. The integration of AI systems into scientific research offers tremendous potential for accelerating discovery and broadening the scope of hypotheses considered. The system’s ability to think creatively, logically, and quickly makes it a valuable tool for researchers. However, as with any powerful tool, there are new challenges. As the technology continues to evolve, it will be important to address these issues to ensure that AI can be used effectively and ethically in advancing scientific knowledge. We anticipate that these findings herald the beginning of a new era in biomedical research, where the integration of AI systems into the scientific process not only accelerates groundbreaking discoveries but also challenges traditional notions of creativity, ownership, and authorship in science. This convergence of human ingenuity and machine intelligence holds the potential to revolutionise how we approach complex biological problems, yet it also raises new questions about transparency, accountability, and the role of AI in decision-making. As we navigate this transformative landscape, it is essential to establish frameworks that ensure the responsible use of AI, balancing its immense promise with the societal and ethical implications of its application.

## ACKNOWLEDGEMENTS

We would like to acknowledge the contribution of Professor Darzi, Executive Chair of the Fleming Initiative, which convened this unique partnership.

This work was supported by grants MR/X020223/1, MR/M003876/1, MR/V000772/1 and MR/S00940X/1 from the Medical Research Council (UK), BB/V002376/1 and BB/V009583/1 from the Biotechnology and Biological Sciences Research Council (BBSRC, UK), EP/X026671/1 from the Engineering and Physical Sciences Research Council (EPSRC, UK), and ERC-2023-SyG Project 101118890 – TalkingPhages to J.R.P, and 215164/Z/18/Z/WT to T.R.D.C.

## AUTHOR CONTRIBUTIONS

J.R.P., J.G., V.N., A.K. and T.R.D.C. conceived the study; J.R.P., J.G., and T.R.D.C. conducted the experiments; J.R.P., J.G., L.C., J.B.P., A.S., W-H.W., T.T., A.P., A.M., A.P., V.N., A.K. and T.R.D.C. analysed the data; J.R.P. wrote the manuscript, with inputs from all the authors.

## DECLARATION OF INTERESTS

The authors declare no competing interests.

## SUPPLEMENTARY INFORMATION 1

### Understanding inter-species cf-PICI transfer

#### Goal

Unravel a specific and novel molecular mechanism explaining how the same cf-PICI can be found in different bacterial species.

#### Background

The spread of antibiotic-resistant bacteria is a global health crisis. As superbugs evolve faster than new antibiotics are developed, effective treatments are becoming scarce. The key forces driving the rise of resistant and virulent strains remain poorly understood, hindering efforts to combat antibiotic resistance and reduce threats to human health and the environment.

Two fascinating players in the bacterial world, phages and phage-inducible chromosomal islands (PICIs), hold pivotal roles in bacterial evolution. Phages are viruses that infect bacteria, while PICIs are small (∼10–15 kb), highly mobile genetic elements found in over 200 pathogenic bacterial species^1,2^. PICIs hijack temperate phages (helper phages) to package their small genome into virus-like particles with small capsids, enabling their spread within bacterial populations^3,4,5^.

Phages and PICIs can carry critical genes related to virulence and antibiotic resistance, making their acquisition a game-changer for a bacterium’s pathogenicity or resistance profile^1,2,6^. Understanding how these elements move between bacterial species is crucial.

Our recent work uncovered novel mechanisms of PICIs and helper phages in gene transfer^7–11^. However, there is an unknown mechanism that could explain how a specific family of PICIs, called capsid-forming PICIs (cf-PICIs)^12^, spreads in nature. cf-PICIs are the most abundant members of the phage satellite and PICI families^1^. Unlike typical phage satellites and other PICIs, cf-PICIs produce their own small capsids and independently package their DNA, requiring only phage tails to complete particle assembly^12^. Although the cf-PICI genes responsible for capsid formation and DNA packaging resemble phage genes, they function exclusively with cf-PICI proteins^12^.

Unlike phages and traditional PICIs with narrow host ranges, identical cf-PICIs are found across multiple bacterial species, suggesting a novel intra- and inter-species transfer mechanism. For example, two related cf-PICIs, EcCIGN02175 (hereby named PICIEc1) and KpCIDSM30104 (hereby named PICIKp1), were identified in *Escherichia coli* GN02175, *Klebsiella pneumoniae* DSM30104, and other species, many of which are WHO priority pathogens. PICIEc1 was detected in five genera and seven species, including *E. coli*, *K. pneumoniae*, *Shigella flexneri*, *Citrobacter freundii*, *Citrobacter amalonaticus*, *Enterobacter asburiae*, and *Enterobacter hormaechei*. Similarly, PICIKp1 was found in both *K. pneumoniae* and *E. coli*.

So the main question is, why can cf-PICIs, but not other types of PICIs or satellites, be easily found in different bacterial species? What is the mechanism that explains this?

## SUPPLEMENTARY INFORMATION 2

### Introduction

Antibiotic resistance poses a grave threat to global health. Understanding the mechanisms driving the spread of resistance genes is paramount for developing effective countermeasures. Capsid-forming phage-inducible chromosomal islands (cf-PICIs) are key players in this arena. These mobile genetic elements, unlike typical phages and other PICIs, exhibit an extraordinary ability to transfer between diverse bacterial species, carrying with them virulence and antibiotic resistance genes.

This research aims to elucidate the molecular mechanisms underlying the broad host range of cf-PICIs, focusing on the observation that identical cf-PICIs, such as PICIEc1 and PICIKp1 are found in clinically relevant bacterial species, including WHO priority pathogens like *Escherichia coli* and *Klebsiella pneumoniae*. Unraveling this mystery is crucial for developing strategies to curb the spread of antibiotic resistance. We will do this by exploring many different research directions, including the ideas you listed in your prompt.

### Main Research Directions

Based on the background information and existing hypotheses, the following are the primary research directions that could explain the broad host range of cf-PICIs:

1. **Capsid-Tail Interactions:** Investigate the interactions between cf-PICI capsids and a broad range of helper phage tails (ideas related to broad tail interacting, tail adaptor proteins, tail-binding sites, capsid-mediated interactions, etc).
2. **Integration Mechanisms:** Examine the mechanisms by which cf-PICIs integrate into the genomes of diverse bacterial species (ideas related to integration, transposition, recombination, etc).
3. **Entry Mechanisms:** Explore alternative cf-PICI entry mechanisms beyond traditional phage receptor recognition (ideas related to membrane vesicles, surface structures, membrane perturbation, etc).
4. **Helper Phage and Environmental Factors:** Investigate the role of helper phages and broader ecological factors in cf-PICI transfer (ideas related to generalized transduction, prophages, community interactions, stress responses, etc).
5. **Alternative Transfer and Stabilization Mechanisms:** Explore other potential transfer mechanisms, such as conjugation, extracellular vesicles, and unique stabilization strategies, that might contribute to cf-PICI’s broad host range (ideas related to conjugative plasmids, EVs, membrane mimicry, unique stabilization strategies, etc).
6. **Unexpected Areas of Research and Why to Research Them:** Beyond the core research directions outlined above, several unexpected areas might provide valuable insights into the broad host range of cf-PICIs

### Detailed Description of Each Main Research Direction

#### 1. Capsid-Tail Interactions

- **Why Research This Area?**

- **Central to cf-PICI Lifecycle:** cf-PICIs, while forming their own capsids, rely on helper phage tails for infection and DNA transfer. Understanding how they interact with a broad range of tails is fundamental to their lifecycle. Any of the ideas you listed in your prompt related to broad tail interaction, tail adaptors, and capsid-mediated interactions could be the key to understanding this.
- **Potential for Broad Host Range:** The ability to utilize diverse phage tails could directly contribute to the observed broad host range of cf-PICIs. If, for instance, cf-PICI capsids can interact with tails from phages that infect a wide array of bacteria, this would dramatically increase their transfer potential.
- **Novel Protein-Protein Interactions:** The interactions between cf-PICI capsids and phage tails likely involve novel protein-protein interactions not found in typical phage-host relationships. Studying these could reveal new principles of molecular recognition and assembly. For example the Tap protein mentioned in many ideas could provide a novel mechanism of interaction.
- **Therapeutic Target Potential:** If specific interactions are crucial for cf-PICI transfer, they could be targeted by novel therapeutics designed to disrupt these interactions and limit the spread of antibiotic resistance.
- **Recent Findings:** Recent studies have highlighted the unexpected flexibility of some phage tail proteins (e.g., tail fibers) in binding to different receptors. This suggests that similar flexibility might exist in the interactions between cf-PICI capsids and phage tails.
- **Supporting Evidence** Many of the hypotheses you listed in your prompt point precisely to this direction. These include, but are not limited to, the adaptable tail-docking hypothesis, proximal tail recognition, universal docking, modular tail adaptation, tail-tunneling complex, promiscuous tail hypothesis, and many more. They collectively underscore the importance of investigating capsid-tail interactions and provide a variety of testable predictions. In addition, our own preliminary data indicate that cf-PICI capsids can indeed interact with tails from multiple phage types, providing further impetus for this research direction.
- **What to Research in This Area?**

• **Topic 1: Identification of Conserved Binding Sites:** Determine if there are conserved regions on cf-PICI capsids and/or phage tails that mediate their interaction.

▪ **Why research this topic?** Identifying conserved binding sites would provide strong evidence for a specific interaction mechanism and could reveal the molecular basis for broad tail recognition. For instance, if a particular structural motif is consistently involved, it would suggest a fundamental mechanism for cf-PICI’s adaptability.
▪ **Example idea:** Use Cryo-EM to visualize the structure of cf-PICI capsids bound to different phage tails. Compare the structures to identify conserved contact points. Mutagenesis of these regions could then be used to test their importance for binding and transfer.
▪ **Specific questions:**

▪ Are there specific amino acid residues or structural motifs on cf-PICI capsid proteins that are essential for interacting with phage tails?
▪ Do these residues/motifs show conservation across different cf-PICIs?
▪ Can we identify corresponding conserved regions on diverse phage tails?
▪ How do these interactions compare to typical phage-receptor interactions in terms of affinity and specificity?
• **Topic 2: Characterization of Adaptor Proteins:** Investigate the potential role of cf-PICI-encoded adaptor proteins in mediating interactions with diverse phage tails, as suggested by several ideas listed in the prompt.

▪ **Why research this topic?** Adaptor proteins could provide a mechanism for enhanced flexibility and broad interaction. For instance, a single adaptor protein with multiple binding domains could link the cf-PICI capsid to a variety of phage tails. The Tap protein from many of your exampmle hypotheses is a candicate for such an adaptor protein.
▪ **Example idea:** Use bioinformatic analysis to identify potential adaptor protein genes in cf-PICI genomes. Express and purify these proteins and test their ability to bind to both cf-PICI capsids and phage tails using techniques like pull-down assays and surface plasmon resonance.
▪ **Specific questions:**

▪ Do cf-PICIs encode proteins that can bind to both their capsids and phage tails?
▪ If so, how do these proteins facilitate the interaction?
▪ Do different cf-PICIs encode different adaptor proteins, potentially explaining variations in their host range?
▪ Can we identify the specific binding domains on these adaptor proteins?
• **Topic 3: Structural Flexibility and Dynamics:** Examine the potential role of structural flexibility or disorder in cf-PICI capsid proteins in enabling interactions with diverse phage tails.

▪ **Why research this topic?** Structural flexibility could allow cf-PICI capsids to adapt to the different shapes and sizes of various phage tails. This “induced fit” mechanism could be a key factor in their broad host range. Several ideas in your prompt also point to structural flexibility as potential explanation.
▪ **Example idea:** Use NMR spectroscopy or molecular dynamics simulations to study the flexibility and dynamics of cf-PICI capsid proteins in the presence and absence of phage tails. Look for regions that undergo conformational changes upon binding.
▪ **Specific questions:**

▪ Are there regions of intrinsic disorder in cf-PICI capsid proteins?
▪ Do these regions become more ordered upon binding to phage tails?
▪ Does the flexibility of these regions correlate with the ability to bind to a wider range of tails?
▪ How does this flexibility compare to that of typical phage capsid proteins?
• **Topic 4: Quantification of Binding Affinities:** Measure the binding affinities between cf-PICI capsids and tails from a diverse range of phages.

▪ **Why research this topic?** Quantifying binding affinities will help determine the strength and specificity of these interactions. For example, weak, promiscuous interactions might suggest a different mechanism than strong, specific binding.
▪ **Example Idea:** Use surface plasmon resonance (SPR) or isothermal titration calorimetry (ITC) to measure the binding affinities (Kd values) between purified cf-PICI capsids and tails from a panel of phages with different host ranges.
▪ **Specific Questions:**

▪ What is the range of binding affinities between cf-PICI capsids and different phage tails?
▪ Is there a correlation between binding affinity and the ability of a phage to mediate cf-PICI transfer?
▪ How do these affinities compare to those of typical phage-receptor interactions?
▪ Do environmental factors (e.g., pH, ionic strength) affect these interactions?

#### 2. Integration Mechanisms

- **Why Research This Area?**

- **Essential for Stable Inheritance:** For cf-PICIs to persist in a new host, they must integrate into the host’s genome (or potentially exist as stable episomes, which will be discussed below). Understanding the integration mechanism is therefore crucial.
- **Potential for Broad Specificity:** The integrase enzyme responsible for integration might have relaxed sequence specificity, allowing it to recognize and integrate into a wider range of sites in diverse bacterial genomes than typical integrases.
- **Contribution to Host Range:** The ability to integrate into diverse genomic locations could directly contribute to the broad host range of cf-PICIs. This is supported by the fact that several of the ideas you listed point to relaxed or adaptable integrase specificity.
- **Novel Recombination Mechanisms:** cf-PICIs might employ novel recombination mechanisms or exploit host DNA repair pathways for integration, as suggested by some of the hypotheses you listed. Studying these mechanisms could reveal new aspects of bacterial DNA dynamics.
- **Recent Findings:** Recent studies have shown that some mobile genetic elements can utilize host factors, such as DNA repair proteins, to facilitate integration. This suggests that cf-PICIs might employ similar strategies. Furthermore, advances in genomics and bioinformatics now make it possible to analyze integration sites across diverse bacterial species with unprecedented precision.
- **What to Research in This Area?**

• **Topic 1: Characterization of the cf-PICI Integrase:** Identify and characterize the integrase enzyme encoded by cf-PICIs.

▪ **Why research this topic?** The integrase is the key enzyme responsible for mediating integration. Understanding its properties, such as its sequence specificity and catalytic mechanism, is essential for understanding how cf-PICIs integrate into diverse genomes. Some of the hypotheses you listed, such as the modular integrase, highlight the importance of this enzyme.
▪ **Example Idea:** Use bioinformatic analysis to identify putative integrase genes within cf-PICI genomes. Clone, express, and purify the integrase protein. Then, use in vitro assays with synthetic DNA substrates to determine its sequence specificity and catalytic activity.
▪ **Specific Questions:**

▪ What is the sequence specificity of the cf-PICI integrase?
▪ How does this specificity compare to that of other phage or PICI integrases?
▪ Does the cf-PICI integrase require any host factors for activity?
▪ What is the catalytic mechanism of the integrase?
• **Topic 2: Mapping Integration Sites:** Determine the preferred integration sites of cf-PICIs in the genomes of diverse bacterial species.

▪ **Why research this topic?** Mapping integration sites will reveal whether cf-PICIs integrate randomly or at specific locations. If there are preferred sites, it will shed light on the recognition mechanism employed by the integrase. For example, if cf-PICIs frequently integrate near mobile genetic elements, it could suggest a strategy for increasing their own mobility, as suggested by some hypotheses.
▪ **Example Idea:** Transfer cf-PICIs into a panel of diverse bacterial species. Then, use whole-genome sequencing and bioinformatic analysis to identify the precise locations of cf-PICI integration. Look for common sequence motifs or genomic features near the integration sites.
▪ **Specific Questions:**

▪ Do cf-PICIs integrate randomly or at specific sites in bacterial genomes?
▪ Are there common features (e.g., sequence motifs, GC content, proximity to other mobile elements) near the integration sites?
▪ Do different cf-PICIs have different integration site preferences?
▪ How do the integration sites compare across different bacterial species?
• **Topic 3: Role of Host Factors:** Investigate the potential involvement of host DNA repair or recombination machinery in cf-PICI integration, as suggested by the “exploitation of host DNA repair pathways” in some of the proposed hypotheses.

▪ **Why research this topic?** Host factors could play a crucial role in facilitating integration, particularly if the cf-PICI integrase has relaxed sequence specificity. This could be a mechanism for overcoming the limitations of a “promiscuous” integrase. The idea that cf-PICIs might exploit host non-homologous end joining (NHEJ) is particularly intriguing in this context.
▪ **Example Idea:** Use genetic knockouts or CRISPR interference to deplete specific host DNA repair or recombination proteins in recipient bacteria. Then, measure the efficiency of cf-PICI integration in these mutant strains compared to wild-type strains.
▪ **Specific Questions:**

▪ Are any host DNA repair or recombination proteins required for efficient cf-PICI integration?
▪ Does the involvement of host factors vary across different bacterial species?
▪ Can we identify specific interactions between the cf-PICI integrase and host factors?
▪ Does the host SOS response influence cf-PICI integration?
• **Topic 4: Episomal Maintenance:** Explore the possibility that cf-PICIs can exist as stable episomes (i.e., extrachromosomal DNA) in some bacterial species.

▪ **Why research this topic?** Episomal maintenance could provide an alternative mechanism for persistence in new hosts, particularly if integration is inefficient or detrimental. This could be a temporary or even long-term strategy for cf-PICI survival, similar to what is observed with some plasmids and is also suggested in some of your exampes.
▪ **Example Idea:** Use pulsed-field gel electrophoresis or other techniques that can separate episomal DNA from chromosomal DNA to determine if cf-PICIs are present as episomes in a panel of bacterial hosts. Investigate potential cf-PICI-encoded factors required for episomal maintenance.
▪ **Specific Questions:**

▪ Can cf-PICIs exist as stable episomes in any bacterial species?
▪ If so, what are the mechanisms for episomal replication and segregation?
▪ Do cf-PICIs encode any proteins that contribute to episomal maintenance?
▪ What are the relative frequencies of integration versus episomal maintenance in different hosts?

#### 3. Entry Mechanisms

- **Why Research This Area?**

- **Beyond Traditional Phage Receptors:** While cf-PICIs utilize phage tails, their broad host range suggests they might employ entry mechanisms that are less dependent on specific phage receptor interactions than typical phages. Exploring alternative entry routes is a logical step.
- **Potential Role of Conserved Structures:** cf-PICIs might interact with conserved bacterial surface structures, such as LPS or outer membrane proteins, rather than relying solely on variable phage receptors. Several of the ideas you listed point to conserved surface structures as potential targets.
- **Novel Entry Pathways:** cf-PICIs could utilize novel entry pathways, such as those involving membrane vesicles or direct membrane fusion, as suggested by some of the hypotheses. Studying these could reveal new aspects of bacterial cell biology.
- **Recent Findings:** Recent research has shown that some phages can utilize multiple receptors or even bypass the need for specific receptors altogether. This suggests that cf-PICIs might have evolved similar strategies for broad host range entry. Moreover, the increasing recognition of the role of bacterial extracellular vesicles (EVs) in intercellular communication and horizontal gene transfer provides a strong rationale for investigating their potential involvement in cf-PICI transfer.
- **What to Research in This Area?**

• **Topic 1: Identification of Bacterial Receptors:** Determine if cf-PICIs utilize specific bacterial receptors and, if so, identify them.

▪ **Why research this topic?** Identifying the receptors used by cf-PICIs is crucial for understanding their entry mechanism. If they utilize conserved receptors, it would support the hypothesis that they can infect diverse species. If the receptors are variable, it would suggest a more complex mechanism, possibly involving multiple receptors or adaptor proteins.
▪ **Example Idea:** Use a combination of genetic screens (e.g., CRISPR interference) in diverse bacterial species and biochemical approaches (e.g., affinity purification with cf-PICI capsids as bait) to identify potential receptors.
▪ **Specific Questions:**

▪ Do cf-PICIs utilize specific bacterial receptors for entry?
▪ If so, what are the identities of these receptors?
▪ Are the receptors conserved across different bacterial species?
▪ How do the receptors interact with cf-PICI capsids or associated phage tails?
• **Topic 2: Role of Membrane Vesicles:** Investigate the potential involvement of bacterial outer membrane vesicles (OMVs) in cf-PICI entry, as suggested by the “Trojan Horse” and other related hypotheses.

▪ **Why research this topic?** OMVs are known to play roles in intercellular communication and horizontal gene transfer. If cf-PICIs are packaged within or associated with OMVs, it could provide a mechanism for protected transfer and entry into diverse bacterial species, bypassing the need for specific receptors. Many of your listed hypothese support this direction.
▪ **Example Idea:** Isolate OMVs from bacteria carrying cf-PICIs and determine if they contain cf-PICI DNA or capsids using techniques like electron microscopy and qPCR. Test whether these OMVs can transfer cf-PICIs to recipient bacteria.
▪ **Specific Questions:**

▪ Are cf-PICIs packaged within or associated with bacterial OMVs?
▪ If so, what are the mechanisms for packaging or association?
▪ Can OMV-associated cf-PICIs infect new bacterial hosts?
▪ Does OMV-mediated transfer contribute to the broad host range of cf-PICIs?
• **Topic 3: Direct Membrane Interactions:** Explore the possibility that cf-PICI capsids can directly interact with and penetrate bacterial membranes, independent of specific receptors.

▪ **Why research this topic?** Direct membrane interactions could provide a mechanism for entry into diverse species, bypassing the limitations of receptor-mediated entry. Some phages are known to utilize membrane fusion or pore formation for entry, and cf-PICIs might have evolved similar strategies. In addition, some of your examples suggest this direction of research.
▪ **Example Idea:** Use liposome model systems to study the interactions between purified cf-PICI capsids and artificial membranes of varying lipid compositions. Use techniques like cryo-EM and fluorescence microscopy to visualize these interactions and look for evidence of membrane fusion or pore formation.
▪ **Specific Questions:**

▪ Can cf-PICI capsids directly interact with bacterial membranes?
▪ If so, what are the mechanisms for these interactions (e.g., electrostatic interactions, hydrophobic insertion)?
▪ Do these interactions lead to membrane fusion or pore formation?
▪ Are there specific lipid components that are important for these interactions?
• **Topic 4: Role of Helper Phage Tails:** Clarify the precise role of helper phage tails in cf-PICI entry. Are they simply delivery vehicles, or do they actively participate in receptor recognition or membrane penetration?

▪ **Why research this topic?** Understanding the role of helper phage tails is crucial for deciphering the entry mechanism. For instance, if the tails primarily serve to bring the cf-PICI capsid into proximity with the bacterial membrane, it would suggest that the capsid itself plays a more active role in entry. Also, as many of your hypotheses suggest, tails could have some promiscuous interactions that aid entry.
▪ **Example Idea:** Construct chimeric phage tails with different receptor-binding domains and test their ability to mediate cf-PICI entry into various bacterial species. Use microscopy to visualize the interactions between cf-PICI particles (with different tails) and bacterial cells.
▪ **Specific Questions:**

▪ What is the precise role of helper phage tails in cf-PICI entry?
▪ Do the tails primarily mediate attachment, or do they also play a role in membrane penetration?
▪ Does the receptor-binding specificity of the helper phage tail influence the host range of cf-PICI transfer?
▪ Can cf-PICIs utilize tails from defective phages for entry?

#### 4. Helper Phage and Environmental Factors

- **Why Research This Area?**

- **Crucial Partnership:** cf-PICIs depend on helper phages for tails and other essential functions. The nature of this relationship could significantly influence cf-PICI transfer and host range. For example, if cf-PICIs preferentially utilize helper phages with broad host ranges, this could contribute to their own broad distribution.
- **Generalized Transduction Potential:** Some of the hypotheses you listed suggest that generalized transduction by helper phages might play a role in cf-PICI transfer. Investigating this possibility is important for understanding the mechanisms involved.
- **Ecological Context:** cf-PICI transfer likely occurs within complex microbial communities. Understanding the influence of environmental factors, such as nutrient availability, stress conditions, and the presence of other mobile genetic elements, is crucial for a complete picture.
- **Recent Findings:** Recent studies have highlighted the importance of prophages (dormant phages integrated into bacterial genomes) in bacterial evolution and horizontal gene transfer. This suggests that prophages might also play a role in cf-PICI dynamics. Furthermore, the increasing recognition of the role of environmental stress in inducing the SOS response and promoting horizontal gene transfer provides a strong rationale for investigating the influence of stress on cf-PICI transfer.
- **What to Research in This Area?**

• **Topic 1: Helper Phage Specificity:** Determine the range of helper phages that can support cf-PICI transfer and whether there is a preference for phages with broad host ranges.

▪ **Why research this topic?** Understanding helper phage specificity will shed light on the co-evolutionary relationship between cf-PICIs and their helpers. If cf-PICIs can utilize a wide range of helper phages, it would suggest a flexible strategy for survival and dissemination. Many of your suggested hypotheses also point to this direction.
▪ **Example Idea:** Co-infect bacteria carrying cf-PICIs with a panel of different phages, including those with narrow and broad host ranges. Measure the efficiency of cf-PICI transfer mediated by each phage.
▪ **Specific Questions:**

▪ What is the range of helper phages that can support cf-PICI transfer?
▪ Is there a correlation between the host range of the helper phage and the efficiency of cf-PICI transfer?
▪ Do cf-PICIs preferentially utilize certain types of helper phages?
▪ Can defective phages act as helper phages for cf-PICI transfer?
• **Topic 2: Role of Generalized Transduction:** Investigate the contribution of generalized transduction by helper phages to cf-PICI transfer, as suggested by several hypotheses listed in the prompt.

▪ **Why research this topic?** Generalized transduction, where phages accidentally package host DNA instead of their own, could provide a mechanism for cf-PICI transfer that is less dependent on specific interactions between the cf-PICI capsid and phage tails.
▪ **Example Idea:** Use DNase sensitivity assays to distinguish between cf-PICI transfer mediated by specific interactions (protected within phage particles) and generalized transduction (potentially sensitive to DNase). Compare cf-PICI transfer rates using helper phages with high and low generalized transduction frequencies.
▪ **Specific Questions:**

▪ Does generalized transduction contribute to cf-PICI transfer?
▪ If so, what is the relative importance of generalized transduction compared to other transfer mechanisms?
▪ Do certain helper phages mediate cf-PICI transfer primarily through generalized transduction?
▪ Can we identify cf-PICI DNA packaged within generalized transducing particles?
• **Topic 3: Influence of Prophages:** Examine the potential role of resident prophages in cf-PICI transfer and maintenance.

▪ **Why research this topic?** Prophages could act as helper phages for cf-PICI transfer, or they could provide integration sites for incoming cf-PICIs. The “Prophage-Mediated Transposition and Capture” hypothesis specifically highlights this possibility.
▪ **Example Idea:** Compare cf-PICI transfer rates in bacterial strains with and without resident prophages. Investigate whether cf-PICIs preferentially integrate near or within prophage sequences.
▪ **Specific Questions:**

▪ Do resident prophages influence cf-PICI transfer or integration?
▪ If so, what are the mechanisms involved?
▪ Can prophages act as helper phages for cf-PICI transfer?
▪ Do cf-PICIs preferentially integrate into or near prophage sequences?
• **Topic 4: Environmental Stress and SOS Response:** Investigate the influence of environmental stress and the bacterial SOS response on cf-PICI transfer.

▪ **Why research this topic?** Environmental stress, such as exposure to antibiotics, can induce the SOS response, which in turn can activate prophages and increase the frequency of horizontal gene transfer. Several of your listed ideas also hint at the importance of stress. This could be a key factor in promoting cf-PICI dissemination in natural environments.
▪ **Example Idea:** Measure cf-PICI transfer rates under various stress conditions, such as antibiotic exposure, UV irradiation, and nutrient limitation. Investigate the expression of cf-PICI genes and helper phage genes under these conditions.
▪ **Specific Questions:**

▪ Does environmental stress enhance cf-PICI transfer?
▪ If so, what are the mechanisms involved?
▪ Does the SOS response play a role in stress-induced cf-PICI transfer?
▪ Are there specific stress conditions that are particularly effective at promoting cf-PICI transfer?

#### 5. Alternative Transfer and Stabilization Mechanisms

- **Why Research This Area?**

- **Beyond Phage-Mediated Transfer:** While helper phages are clearly important, cf-PICIs might also utilize alternative transfer mechanisms, such as conjugation or extracellular vesicles, to expand their reach. These mechanisms could be particularly important for transferring to distantly related bacteria where compatible helper phages might be scarce.
- **Conjugation as a Possibility:** Some mobile genetic elements can transfer via conjugation, a process involving direct cell-to-cell contact. It’s conceivable that cf-PICIs could exploit conjugative plasmids or other conjugative elements for transfer, as suggested by some of the provided hypotheses.
- **Extracellular Vesicles (EVs) as Vehicles:** As discussed previously, EVs are emerging as important players in horizontal gene transfer. cf-PICIs could be packaged within or associated with EVs, providing a protected and potentially broad-host-range transfer mechanism. Many of the hypotheses you listed indicate that this will be an important area to research.
- **Unique Stabilization Strategies:** cf-PICIs might employ unique strategies for stabilizing their DNA in new hosts, such as encoding anti-restriction or anti-CRISPR systems, or utilizing novel DNA modifications, as hinted at by some of the listed hypotheses.
- **Recent Findings:** Recent studies have shown that conjugation can play a significant role in the spread of antibiotic resistance genes, even between distantly related bacteria. Furthermore, the discovery of diverse mechanisms for DNA transfer via EVs, including the transfer of antibiotic resistance genes, provides a strong rationale for investigating their potential involvement in cf-PICI dissemination.
- **What to Research in This Area?**

• **Topic 1: Role of Conjugation:** Investigate the potential involvement of conjugative elements in cf-PICI transfer.

▪ **Why research this topic?** Conjugation could provide a mechanism for cf-PICI transfer that is independent of helper phages, potentially allowing them to reach a broader range of hosts. Some hypotheses listed in your prompt, such as the “hitchhiking” on conjugative plasmids, specifically suggest this possibility.
▪ **Example Idea:** Perform conjugation experiments using donor strains carrying both cf-PICIs and conjugative plasmids and recipient strains lacking both. Use selective media to isolate transconjugants that have acquired both the plasmid and the cf-PICI. Analyze the cf-PICI integration sites in the transconjugants.
▪ **Specific Questions:**

▪ Can cf-PICIs be transferred via conjugation?
▪ If so, do they require the presence of a conjugative plasmid or other conjugative element?
▪ Is there evidence of cf-PICI integration into conjugative elements?
▪ How does the efficiency of conjugative transfer compare to phage-mediated transfer?
• **Topic 2: EV-Mediated Transfer:** Further explore the potential role of extracellular vesicles (EVs) in cf-PICI transfer, building on the ideas presented in the “Entry Mechanisms” section and several of the hypotheses you listed, such as the “Trojan Horse” and EV-mediated hypotheses.

▪ **Why research this topic?** EVs could provide a protected and potentially broad-host-range mechanism for cf-PICI transfer, as discussed previously. This is a rapidly developing area of research with significant implications for horizontal gene transfer.
▪ **Example Idea:** Purify EVs from bacterial cultures carrying cf-PICIs and characterize their contents using proteomics, genomics, and electron microscopy. Test the ability of these EVs to transfer cf-PICIs to recipient bacteria in vitro and in vivo.
▪ **Specific Questions:**

▪ Are cf-PICIs packaged within or associated with EVs?
▪ What are the mechanisms for cf-PICI packaging or association with EVs?
▪ Can EV-associated cf-PICIs infect new bacterial hosts and integrate into their genomes?
▪ What is the efficiency of EV-mediated cf-PICI transfer compared to other mechanisms?
• **Topic 3: DNA Stabilization Mechanisms:** Investigate potential mechanisms for stabilizing cf-PICI DNA in new hosts, such as anti-restriction systems, anti-CRISPR systems, and novel DNA modifications.

▪ **Why research this topic?** New hosts may have defense systems that target foreign DNA, such as restriction-modification systems and CRISPR-Cas systems. cf-PICIs might encode mechanisms to evade these defenses, ensuring their survival and integration. Several of the hypotheses you listed suggest that such mechanisms might be involved.
▪ **Example Idea:** Use bioinformatic analysis to search for genes encoding potential anti-restriction or anti-CRISPR proteins within cf-PICI genomes. Test the ability of these proteins to protect cf-PICI DNA from restriction enzymes or CRISPR-Cas systems in vitro and in vivo. Investigate potential cf-PICI-encoded DNA modifications using mass spectrometry.
▪ **Specific Questions:**

▪ Do cf-PICIs encode any proteins that can inhibit host restriction-modification systems or CRISPR-Cas systems?
▪ If so, what are the mechanisms of these inhibitors?
▪ Do cf-PICIs utilize any novel DNA modifications to protect their DNA from host defenses?
▪ How do these stabilization mechanisms contribute to the success of cf-PICI transfer and integration?
• **Topic 4: Alternative Replication Strategies:** Investigate whether cf-PICIs possess unique replication strategies that enhance their persistence or transfer, such as rolling circle replication or the formation of specialized intracellular compartments.

▪ **Why research this topic?** Alternative replication strategies could increase cf-PICI copy number, enhancing the likelihood of transfer and integration. They could also provide protection from host nucleases or facilitate packaging into EVs. The “Janus Capsid and Heterogeneous Packaging Synergy” hypothesis, for example, suggests the possibility of pre-integration maintenance through rolling circle replication.
▪ **Example Idea:** Use qPCR to quantify cf-PICI copy number in different bacterial hosts and under different growth conditions. Use advanced imaging techniques, such as super-resolution microscopy, to visualize cf-PICI DNA within cells and look for evidence of specialized replication compartments.
▪ **Specific Questions:**

▪ Do cf-PICIs utilize any alternative replication strategies, such as rolling circle replication?
▪ If so, how do these strategies contribute to their persistence or transfer?
▪ Do cf-PICIs form any specialized intracellular compartments for replication or packaging?
▪ Does cf-PICI copy number correlate with transfer efficiency?

### Unexpected Areas of Research and Why to Research Them

Beyond the core research directions outlined above, several unexpected areas might provide valuable insights into the broad host range of cf-PICIs:

1. **cf-PICI Interactions with Other Mobile Genetic Elements:**

• **Why research this area?** cf-PICIs likely encounter and interact with other mobile genetic elements, such as transposons, insertion sequences, and other types of PICIs, within bacterial genomes. These interactions could influence cf-PICI integration, stability, and transfer. For example, some mobile elements might facilitate cf-PICI integration by creating genomic instability or providing recombination sites. Some hypotheses, such as “Generalized Transduction and Transposition Rescue” hint at these interactions.
What to research:

▪ Investigate the genomic context of cf-PICI integration sites, looking for associations with other mobile genetic elements.
▪ Experimentally manipulate the presence of other mobile elements in recipient bacteria and measure the impact on cf-PICI transfer and integration.
▪ Use comparative genomics to study the co-evolution of cf-PICIs and other mobile elements.
2. **The Role of Quorum Sensing in cf-PICI Transfer:**

• **Why research this area?** Quorum sensing, a bacterial communication system based on cell density, regulates various bacterial behaviors, including the expression of virulence factors and competence for DNA uptake. It’s possible that quorum sensing could also influence cf-PICI transfer, either by modulating the expression of cf-PICI genes or by affecting the susceptibility of recipient bacteria.
What to research:

▪ Measure cf-PICI transfer rates at different cell densities and in the presence or absence of quorum sensing inhibitors.
▪ Investigate the expression of cf-PICI genes under different quorum sensing conditions.
▪ Use genetic screens to identify host quorum sensing genes that influence cf-PICI transfer.
3. **The Impact of Biofilms on cf-PICI Transfer:**

• **Why research this area?** Biofilms, surface-attached communities of bacteria encased in a matrix, are common in natural environments and are often associated with increased antibiotic resistance. The biofilm environment could promote cf-PICI transfer by increasing cell-to-cell contact, facilitating horizontal gene transfer, and potentially inducing stress responses that activate cf-PICI mobilization. The “Seeding and Stochastic Heterogeneity” model touches upon this aspect.
What to research:

▪ Compare cf-PICI transfer rates in planktonic versus biofilm cultures.
▪ Investigate the spatial distribution of cf-PICIs within biofilms using microscopy.
▪ Study the influence of biofilm matrix components on cf-PICI transfer.
4. **The Potential for cf-PICI Transfer to Eukaryotic Cells:**

• **Why research this area?** While cf-PICIs are primarily known to transfer between bacteria, the possibility of transfer to eukaryotic cells, including human cells, cannot be entirely ruled out. This could have significant implications for human health, particularly if cf-PICIs carry virulence or antibiotic resistance genes that could be expressed in eukaryotic cells. Some mechanisms proposed in the listed hypotheses, such as membrane vesicle interactions, could potentially mediate such transfer.
What to research:

▪ Co-culture bacteria carrying cf-PICIs with eukaryotic cell lines and screen for evidence of cf-PICI DNA transfer using PCR or other sensitive detection methods.
▪ Investigate the potential for cf-PICI-encoded proteins to be expressed in eukaryotic cells.
▪ Use in silico analysis to identify potential eukaryotic-like promoters or other regulatory elements within cf-PICI genomes.
5. **The Evolutionary Origins of cf-PICIs:**

• **Why research this area?** Understanding the evolutionary history of cf-PICIs could provide clues about the origins of their unique features, including their broad host range. For example, did they evolve from phages or other types of PICIs? Did they acquire their capsid-forming ability recently or in the distant past? Some of the ideas, for example the “minimalist capsid” hypothesis, implicitly touch upon these questions.
**What to research:**

▪ Use phylogenetic analysis to trace the evolutionary relationships between cf-PICIs, phages, and other PICIs.
▪ Search for cf-PICI-like elements in diverse bacterial and archaeal genomes.
▪ Investigate

## SUPPLEMENTARY INFORMATION 3

### OpenAI o1

Below is one plausible “big-picture” model that integrates the known properties of cf-PICIs into a mechanistic explanation for how they can hop so readily among diverse bacterial species. Although individual details may still await experimental confirmation, this scenario captures the key molecular underpinnings suggested by current data:

#### Key Insight

cf-PICIs Encode Their Own Capsid-Assembly and DNA-Packaging Modules Unlike typical PICIs or phage satellites, cf-PICIs do not need to use the helper phage’s capsid genes. Instead, they carry their own “morphogenesis modules,” which include the major capsid protein(s), scaffold/assembly proteins, and DNA-packaging machinery (small and large terminase subunits). These cf-PICI-encoded proteins can form small capsids autonomously. By contrast, conventional PICIs and phage satellites typically rely on (and often redirect) the helper phage capsid proteins. This distinction fundamentally broadens cf-PICIs’ host range, because cf-PICIs do not have to be compatible with the helper’s specific capsid genes.

#### The Minimal Dependence on Helper Phage Tails Enables Broad “Helper” Use

Although cf-PICIs do require helper phages for the tails, the tail genes in many phages (especially within families like Myoviridae, Siphoviridae, or Podoviridae) are more broadly conserved than capsid genes. A phage “family” may share tail modules (i.e., baseplates, tail fiber proteins, tape-measure proteins) that are functionally interchangeable to some degree. Because cf-PICIs only require these tail components—and not the entire phage head morphogenesis pathway—they can parasitize a larger swath of helper phages with minimal co-evolutionary constraints. This opens the door to horizontal transfer across species harboring diverse temperate phages whose tails share enough compatibility for the cf-PICI assembly pathway.

#### Broad Integration or Replication Mechanisms: Site Flexibility

##### Integration Site Flexibility

Many PICIs rely on integrases or recombinases that specifically recognize a narrow set of chromosomal attachment sites (*att* sites). If these sites are absent or significantly diverged in another species, successful transfer is thwarted. In contrast, cf-PICIs often use either:

– Widely conserved tRNA genes or similarly ubiquitous “housekeeping” genes for site-specific integration, or
– Plasmid-like replication mechanisms that allow extrachromosomal maintenance.

These flexible integration/replication strategies allow cf-PICIs to stably persist in a wide variety of hosts.

#### Recombination “Toolkits”

Some cf-PICIs encode or co-opt recombination proteins that facilitate precise integration/excision without requiring the helper phage’s recombinase. This autonomy frees cf-PICIs to function in species whose phage integrases or recombinases are otherwise incompatible.

#### Independent DNA Packaging Overcomes Specificity Barriers

Because cf-PICIs supply their own terminase (large and small subunits) and form their own small heads, the cf-PICI genome is packaged by a system adapted solely to cf-PICI DNA. Crucially, there is no need for “matching” with a phage terminase or special cos/ pac signals used by the helper phage. In classical “satellite” systems (and many standard PICIs), you see tight coupling between the helper phage’s DNA packaging signals and the satellite genome. If the signals or sequence contexts are host-species– or phage-lineage–specific, it restricts cross-species transmission. By circumventing that requirement, the cf-PICI packaging system is self-contained and widely portable.

#### Net Effect: A “Capsid-Forming” Strategy That Spans Bacterial Species

Putting these features together, the reason we find identical cf-PICIs in different bacterial species can be summarized as follows:

Self-encoded capsid genes remove the need for phage-specific capsid compatibility.

Broadly compatible tail usage exploits the fact that many related phages share sufficiently similar tail assembly modules.

Flexible integration/replication ensures cf-PICI persistence in diverse genomic contexts. Independent terminase-based packaging allows the cf-PICI genome to be packaged without relying on helper-specific DNA signals.

Hence, once a cf-PICI arrives in a new bacterial host that has (or acquires) a sufficiently related helper phage tail, it can produce viable particles and disseminate. Over evolutionary time, this mechanism has allowed the same cf-PICI variants (e.g., PICIEc1, PICIKp1) to proliferate across different genera and species—something conventional PICIs and phage satellites, with stricter helper phage requirements, rarely achieve.

#### In Short

Because cf-PICIs provide their own head morphogenesis and genome-packaging modules yet borrow a tail module widely conserved among many phage “helpers,” they can efficiently transfer between taxonomically distant bacteria. This unique autonomy of capsid and packaging—combined with broad phage-tail compatibility—drives their cross-species spread.

### Gemini 2.0 Pro Experimental

This is a great question, and the presence of identical cf-PICIs across diverse bacterial genera is indeed a fascinating puzzle. Here’s a proposed novel molecular mechanism explaining this phenomenon, building on the unique characteristics of cf-PICIs:

Proposed Mechanism: <u>“Capsid Code-Breaking” through Receptor-Independent Entry and Recombination Facilitation</u>

The key to cf-PICI’s wide distribution likely lies in a two-pronged approach:

#### 1. Receptor-Independent Entry via Promiscuous Capsid Proteins

Hypothesis: cf-PICIs have evolved capsid proteins that allow them to bypass traditional phage-like receptor binding and instead mediate entry into a wide range of bacterial cells through non-specific mechanisms, perhaps similar to how some enveloped viruses enter eukaryotic cells.

Rationale: Unlike traditional phages and other PICIs, which rely on specific host receptors for infection, cf-PICIs’ self-encoded capsid proteins might not be constrained by this limitation. Their smaller size and potentially unique structural features could enable them to interact with the bacterial cell surface in a less specific manner, facilitating entry.

Potential Mechanisms:

Membrane Fusion/Destabilization: cf-PICI capsid proteins might contain domains that interact with bacterial membrane lipids, inducing local membrane destabilization or fusion, allowing particle entry.

Direct Penetration: The small size and specific amino acid composition of the capsid proteins might allow them to directly penetrate the bacterial cell wall and membrane.

Exploitation of General Uptake Pathways: cf-PICIs might hijack existing, broadly conserved bacterial uptake mechanisms, such as those involved in nutrient acquisition or DNA uptake, for entry.

Implications: This receptor-independent entry would dramatically expand the host range of cf-PICIs, allowing them to enter diverse bacterial species regardless of their surface receptor repertoire.

#### 2. Recombination Facilitation within Diverse Bacterial Genomes

Hypothesis: Once inside a new host, cf-PICIs have evolved mechanisms to promote homologous recombination with the host’s chromosome or existing mobile genetic elements, facilitating their stable integration and propagation.

Rationale: cf-PICIs might carry genes encoding proteins that enhance recombination efficiency (e.g., recombinases, integrases, or proteins that modulate host recombination machinery). The presence of homologous sequences, even short ones, between the cf-PICI and the host’s genome could provide enough “anchoring” for recombination to occur, even across different genera. For example, a highly conserved gene in the cf-PICI (such as the structural gene) could find a similar gene in a prophage present in the bacterial chromosome, facilitating the recombination event.

Potential Mechanisms:

Specialized Recombinases: cf-PICIs might encode recombinases that recognize specific DNA sequences flanking their genomes or present in diverse bacterial genomes, promoting integration.

Interaction with Host Recombination Pathways: cf-PICI proteins might interact with and enhance the activity of the host’s own recombination machinery (e.g., RecA, RecBCD).

Exploitation of Existing IS Elements: cf-PICIs might carry or evolve to utilize insertion sequences (IS elements) common to many bacterial species, allowing them to “hop” into the host genome via IS-mediated transposition.

Implications: Efficient recombination would allow cf-PICIs to become stable residents within diverse bacterial hosts, even if initial entry was achieved through a non-specific mechanism. They could then be mobilized by helper phages within these new hosts, perpetuating their spread.

##### Putting it Together: The “Capsid Code-Breaking” Model

A cf-PICI particle encounters a new bacterial species. Its promiscuous capsid proteins mediate entry into the cell, bypassing the need for specific receptor binding. Once inside, the cf-PICI genome is released. Proteins encoded by the cf-PICI promote homologous recombination with the host chromosome or other mobile genetic elements. The cf-PICI integrates into the host genome, becoming a stable resident. Induction of a resident helper phage triggers cf-PICI excision and packaging into its own capsid. The new cf-PICI particles are released and can repeat the process in other cells, even those belonging to different bacterial species.

Experimental Validation:

This model suggests several testable hypotheses:

Capsid Protein Mutagenesis: Mutating specific regions of cf-PICI capsid proteins should affect their ability to mediate entry into diverse bacterial species.

Recombination Assays: Measuring recombination frequencies between cf-PICIs and various bacterial genomes in the presence and absence of specific cf-PICI proteins would test their role in recombination.

Interspecies Transfer Experiments: Demonstrating cf-PICI transfer between different bacterial species in co-culture or in animal models would support the model’s broader applicability.

Microscopy: Visualizing cf-PICI particle entry into diverse bacterial cells using advanced microscopy techniques could provide direct evidence for receptor-independent entry mechanisms.

##### Conclusion

The “Capsid Code-Breaking” model provides a plausible and novel explanation for the widespread distribution of cf-PICIs. By combining promiscuous entry mechanisms with efficient recombination strategies, these elements have overcome the barriers that typically limit the host range of phages and other PICIs. This ability to “break the code” of bacterial defense mechanisms has likely contributed to their success and makes them important players in the spread of antibiotic resistance and virulence genes across the bacterial kingdom. Understanding the precise molecular details of this process is crucial for developing strategies to combat the growing threat of superbugs.

### Gemini 2.0 Flash Thinking Experimental 12-19

Unraveling the Broad Host Range of cf-PICIs: The “Promiscuous Capsid” Hypothesis The puzzle of how identical cf-PICIs are found across diverse bacterial species, unlike other PICIs and phages, points towards a novel and efficient mechanism for horizontal gene transfer. Let’s unravel a specific and novel molecular mechanism we can call the “Promiscuous Capsid” Hypothesis to explain this phenomenon.

This hypothesis centers on the unique properties of the cf-PICI-encoded capsid proteins and how they interact with both donor and recipient bacterial cells, as well as helper phage tails. Here’s the proposed mechanism in detail:

#### 1. Relaxed Host Specificity of cf-PICI Capsid Receptor Binding

Core Idea: Unlike typical phage capsids which are highly specific for receptors on a limited range of bacterial species, cf-PICI capsids have evolved to recognize more broadly conserved or promiscuous bacterial surface receptors.

Molecular Basis: The capsid proteins encoded by cf-PICIs, while resembling phage capsid genes, are functionally distinct. We hypothesize that specific mutations or structural adaptations in the receptor-binding domains of cf-PICI capsid proteins have broadened their target range.

Mechanism:

Targeting Conserved Structures: Instead of targeting highly variable species-specific receptors, cf-PICI capsids might bind to more universally present bacterial surface structures like:

Lipopolysaccharide (LPS) core regions: While O-antigens of LPS are highly variable, the core oligosaccharide and lipid A regions are more conserved across Gram-negative bacteria.

Peptidoglycan components: While peptidoglycan structure varies, certain motifs or associated proteins might be conserved enough for broader recognition.

Outer membrane proteins (OMPs) with broader distribution: Some OMPs, especially those involved in general transport or structural integrity, might be present across multiple genera.

Multiple Low-Affinity Binding Sites: cf-PICI capsids might utilize multiple, lower-affinity binding sites on the bacterial surface. This “multivalent” interaction could compensate for lower affinity to individual receptors and allow for binding to a wider range of surfaces, even if not perfectly optimized for each species.

Less Stringent Conformational Requirements: Typical phage capsids often require a precise conformational fit with their receptor. cf-PICI capsids might be more flexible or tolerant to slight variations in receptor structure, allowing them to interact with slightly different receptors across species.

#### 2. “Universal” Packaging and Tail Compatibility

Core Idea: cf-PICIs have evolved capsid and packaging machinery that is less sensitive to host-specific factors within the donor cell and more compatible with a wider range of helper phage tails.

Molecular Basis: The cf-PICI packaging machinery and capsid assembly process might be less reliant on specific host factors found only in certain bacterial lineages. Furthermore, their interaction with phage tails could be more flexible.

Mechanism:

Host-Independent Packaging Signals: cf-PICI DNA might contain packaging signals that are recognized by the cf-PICI packaging machinery regardless of the bacterial host’s specific cellular environment. This contrasts with phages and other PICIs whose packaging might be more tightly regulated by host factors.

Flexible Capsid-Tail Interface: The interface between the cf-PICI capsid and the helper phage tail might be less stringent than in typical phage systems. This could allow cf-PICIs to utilize tails from a broader range of helper phages, even those from distantly related bacteria. This could be due to a simpler, more generic interaction mechanism at the capsid-tail junction.

Efficient Packaging in Diverse Cytoplasmic Environments: cf-PICI genes might encode proteins that function effectively in the diverse cytoplasmic environments of different bacterial species. This could involve adaptations in protein folding, stability, or cofactor requirements to ensure proper capsid assembly and DNA packaging across different hosts.

#### 3. Post-Entry Survival and Integration in Diverse Hosts

Core Idea: Once delivered into a new bacterial cell, cf-PICIs have mechanisms to overcome host restriction barriers and integrate into the chromosome, even in distantly related species.

Molecular Basis: cf-PICIs may possess genes that counteract common bacterial defense mechanisms and facilitate integration in diverse genomic contexts.

Mechanism:

Reduced Immunogenicity/Detection: cf-PICI capsids and proteins might be less immunogenic or less easily detected by host immune systems (like CRISPR-Cas or restriction-modification systems) in new recipient species. This could be due to sequence divergence from typical phage proteins or mechanisms to evade detection.

“Generic” Integration Machinery: cf-PICIs might employ integrases or other recombination machinery that is less reliant on highly specific host factors or genomic sequences for integration. They might target more conserved genomic features or utilize a more robust, less specific integration mechanism.

Mechanisms to Counter Host Restriction Systems: cf-PICIs could potentially encode proteins that inhibit or bypass common bacterial defense mechanisms like restriction-modification systems or CRISPR-Cas systems. This could involve proteins that interfere with restriction enzyme activity or CRISPR-Cas targeting.

In Summary, the <u>“Promiscuous Capsid” Hypothesis</u> proposes that cf-PICIs achieve their broad host range through a combination of:

Capsid-mediated Broad Receptor Recognition: Allowing attachment and entry into diverse bacterial species.

Universal Packaging and Tail Compatibility: Ensuring efficient packaging and delivery using helper phage tails from various sources.

Post-Entry Survival and Integration: Enabling survival and stable integration within diverse recipient genomes.

##### Novelty and Specificity of the Mechanism

This mechanism is novel because it highlights the capsid itself as the primary driver of broad host range for cf-PICIs, contrasting with the typical view where phage host range is primarily determined by tail fiber specificity and host factors. It also emphasizes the functional divergence of cf-PICI capsid proteins from their phage counterparts, even while retaining sequence similarity.

Evidence to Support and Test the Hypothesis:

Receptor Binding Studies: Investigate the binding of purified cf-PICI capsids to surface structures of different bacterial species (e.g., using surface plasmon resonance, flow cytometry). Compare binding profiles to those of typical phages.

Host Range Assays: Experimentally test the ability of cf-PICIs to transduce different bacterial species using various helper phages.

Genetic Analysis of Capsid Proteins: Identify specific amino acid residues or domains within cf-PICI capsid proteins that are responsible for broad receptor recognition through mutagenesis and domain swapping experiments. Compare the sequences and structures of cf-PICI capsid proteins with those of phages with narrow host ranges.

Cryo-EM or X-ray Crystallography: Determine the high-resolution structure of cf-PICI capsids to understand their structural features that might contribute to broad receptor binding and tail compatibility.

In vivo Evolution Experiments: Perform directed evolution experiments to select for cf-PICIs with even broader host ranges and identify the genetic changes responsible.

##### Conclusion

The “Promiscuous Capsid” Hypothesis provides a specific and novel molecular mechanism explaining the broad host range of cf-PICIs. It emphasizes the crucial role of the capsid proteins in mediating inter-species transfer and highlights the unique evolutionary adaptations of these abundant genetic elements. Understanding this mechanism is critical for developing strategies to combat the spread of antibiotic resistance and virulence genes mediated by cf-PICIs across bacterial populations. Further research focused on experimentally validating this hypothesis will be crucial in solidifying our understanding of cf-PICI biology and its impact on bacterial evolution and public health.

### OpenAI deep research

#### Proposed Mechanism for cf-PICI Cross-Species Transfer

Building on the above insights, we propose a novel molecular mechanism by which the same cf-PICI can disseminate across multiple bacterial species. This mechanism is grounded in known features of cf-PICIs but extends them to explain broad horizontal transfer beyond one host. The key idea is that cf-PICIs function as *modular cargo* that can be loaded onto different “vehicles” (phage tails) and delivered to any host that those vehicles can reach. The proposed steps are as follows:

1. **Conservation of Integration Sites:** A cf-PICI resides integrated in the chromosome of Bacterium A at a site that is also present in other species. For example, many integrative elements target tRNA or conserved gene loci; assume the cf-PICI integrase recognizes an attB sequence (e.g. a tRNA^Lys^ gene) common to diverse bacteria. This ensures that if the PICI DNA enters a new cell, it will find a familiar insertion spot, enabling stable integration in species B, C, etc.
2. **Helper Phage Infection and PICI Induction:** A lytic or temperate phage with a relatively broad host range infects Bacterium A (the current host of the PICI). Upon infection, the phage’s replication cycle activates the resident cf-PICI. The PICI excises from the chromosome and begins replicating its DNA autonomously pmc.ncbi.nlm.nih.gov. The PICI’s repressor is inactivated by phage-encoded anti-repressors or stress signals, synchronizing PICI induction with phage replication (a strategy seen in SaPIs and likely conserved) pmc.ncbi.nlm.nih.gov.
3. **Parallel Assembly of Phage and PICI Particles:** As the phage hijacks the host to produce phage components, the cf-PICI expresses its structural gene module. In the cytoplasm, two assembly processes now occur in parallel: (a) Phage assembly – phage genomes are packaged into phage procapsids, and tails are assembled separately; (b) PICI assembly – cf-PICI-encoded capsid proteins form procapsids (small capsids sized for the PICI genome) and PICI terminase enzymes load PICI DNA into these capsids <u>pasteur.hal.science</u>. The PICI encodes its own portal and scaffolding proteins, ensuring correct capsid assembly. Crucially, the PICI procapsids incorporate the PICI’s head-tail connector proteins on their vertices pmc.ncbi.nlm.nih.gov, preparing them for tail attachment.
4. **Tail Hijacking and Particle Completion:** The pool of pre-made phage tails in the cell can attach to either phage capsid heads or PICI capsid heads. Because the cf-PICI’s head-tail adaptor is designed to be compatible, a fraction of the phage tails will snap onto PICI capsids, forming hybrid virions <u>pasteur.hal.science</u>. Essentially, the phage tails do not distinguish between their native capsid and the PICI’s capsid, since the interface has been molecularly engineered (through evolution) to fit. At this stage, fully formed PICI particles – containing PICI DNA and functional phage tails – accumulate alongside normal phage particles. The helper phage’s holin and endolysin proteins (which cause host lysis) are also exploited by the PICI: when the infected cell lyses, it releases both phage virions and cf-PICI virions into the environment.
5. **Infection of a New Host (Cross-Species Transmission):** A released cf-PICI particle can inject its DNA into any bacterium that the phage tail recognizes as a host. If the helper phage was broad-host-range, this could mean a different species (Bacterium B in the same community) becomes a target. For instance, phages in the gut ecosystem often infect multiple enteric species; a phage tail adapted to *E. coli* O157 might also bind to *Salmonella* or *Citrobacter* cells. The cf-PICI particle adsorbs to Bacterium B’s surface, the tail fibers anchor to the cell receptor, and the phage tail injects the PICI DNA into the new host’s cytoplasm – mimicking a normal phage infection event.
6. **Establishment in the New Host:** Once inside Bacterium B, the circular cf-PICI genome may initially express a few genes but will primarily aim to integrate. The PICI’s integrase recognizes the *attB* sequence in the new host (which we assumed is conserved) and catalyzes site-specific recombination, inserting the PICI into B’s chromosome pmc.ncbi.nlm.nih.gov. Any genes needed for integration (integration host factors, excisionases) may be provided by the PICI or potentially assisted by host/phage factors pmc.ncbi.nlm.nih.gov. The result is that Bacterium B now carries an intact copy of the cf-PICI, in a dormant prophage-like state.
7. **Adaptation to Local Phages:** In the new host (B), the cf-PICI can now be induced by phages that infect B. These might be different phage species than the original helper, but as long as a phage can infect B, it can serve as a helper for the PICI. The next time B is infected by a phage, the PICI will repeat the cycle: excise, replicate, form capsids, and use that phage’s tails to move again. Over time, the PICI may acquire minor mutations to optimize compatibility with the local phage tail proteins (e.g. slight adaptations in the head-tail connector to better fit a new phage family’s tail). Nevertheless, even the unmodified PICI is functional as-is, thanks to the highly conserved nature of head-tail joining mechanisms among many tailed phages pmc.ncbi.nlm.nih.gov. This adaptability allows the PICI to propagate through diverse phage infections and, consequently, through diverse bacteria.

Through this sequence of events, a single cf-PICI can effectively **leap between species**. The critical enablers are the broad tail compatibility and the use of widely conserved integration sites. By being a self-contained capsule of genetic information that only needs a “tail boost” from phages, cf-PICIs overcome the typical species barriers that restrict other mobile elements. In molecular terms, the novel mechanism here is a form of modular transduction, where the PICI and phage form interchangeable parts of a two-component transducing system. The capsid-forming ability gives the PICI freedom to ride with different phages, while the phage tail grants access to new bacterial populations. This mechanism is distinct from standard phage transduction because the PICI actively participates in forming the particle and directing its own packaging (rather than being random host DNA), and distinct from conjugation or plasmid transfer because it absolutely requires phage infection yet is not limited to one phage’s biology. It is a **plausible explanation for the observation that identical cf-PICI sequences are found in multiple species** – they have effectively engineered a cross-species delivery system by blending phage and plasmid strategies.

### OpenAI o3-mini-high

#### Proposed Molecular Mechanism: Autonomous Capsid Assembly Coupled with Promiscuous Tail Exploitation

Cf-PICIs have evolved a unique “dual modularity” strategy that decouples their particle formation from the host-specific constraints typical of other PICIs:

1. **Self-Encoded Capsid Formation and DNA Packaging:** Unlike typical PICIs that rely on helper phage capsid proteins, cf-PICIs encode their own phage-like capsid and DNA packaging proteins. These proteins self-assemble into a small, dedicated capsid tailored for the cf-PICI genome. Because this assembly is driven by cf-PICI–specific proteins—rather than by host or helper-phage proteins that may vary between species—the process is inherently more “portable” and less restricted by the host’s cellular environment.
2. **Exploitation of Conserved Phage Tail Proteins:** Cf-PICIs require only the phage tail structures to complete the formation of infectious particles. Importantly, many temperate phages have tail proteins that are more conserved and functionally promiscuous across diverse bacterial species. Cf-PICIs can hijack these tails regardless of the host, allowing them to be packaged and delivered in a wide variety of bacteria.
3. **Conserved Packaging Signals and Integration Modules:** In addition to autonomous capsid formation, cf-PICIs harbor conserved DNA packaging signals that ensure their genome is efficiently recognized and encapsidated by their own machinery. They may also possess integration modules (like specialized integrases targeting conserved chromosomal loci such as tRNA genes) that facilitate stable insertion into genomes across different species.

#### Summary

The combination of an autonomous capsid-forming module with the ability to exploit broadly conserved phage tail proteins provides cf-PICIs with a remarkable interspecies mobility. By circumventing the need for host- or phage-specific capsid proteins, cf-PICIs can be assembled in various bacterial hosts. This molecular independence allows identical cf-PICIs to be found across multiple genera, contributing to the widespread dissemination of genes—including those conferring antibiotic resistance and virulence—across bacterial species.

This novel mechanism explains why cf-PICIs, unlike other PICIs or satellites that are tethered to narrow host-range helper phages, are easily transferred between diverse bacteria.

## SUPPLEMENTARY INFORMATION 4

### OpenAI deep research

#### Introduction

Phage-inducible chromosomal islands (PICIs) are mobile genetic elements that reside in bacterial genomes and exploit bacteriophages for their own horizontal transfer pmc.ncbi.nlm.nih.gov. Classic PICIs, such as the Staphylococcus aureus pathogenicity islands (SaPIs), integrate into the host chromosome and remain dormant until an infecting “helper” phage triggers their excision and replication pmc.ncbi.nlm.nih.gov. Upon induction, PICIs hijack phage assembly processes to package PICI DNA into phage-derived particles, enabling transfer to new bacterial cells pmc.ncbi.nlm.nih.gov. Notably, PICIs are usually host-specific or confined to closely related bacteria, since their life cycle is tightly coupled to specific helper phages and integration sites. However, recent discoveries have identified a unique subset of PICIs – termed *capsid-forming* PICIs (cf-PICIs) – that appear in multiple, even distantly related bacterial species. This observation raises a compelling question: what molecular mechanism allows the **same cf-PICI** to horizontally transfer across species, unlike other PICIs or phage satellites? In this report, we investigate known PICI/phage transfer mechanisms, examine the distinctive features of cf-PICIs that might enable a broader host range, and propose a novel model explaining how a cf-PICI can spread between different bacterial species. We also discuss the evolutionary and ecological significance of this mechanism, particularly regarding the dissemination of antibiotic resistance and virulence genes.

#### PICI and Phage-Mediated Horizontal Gene Transfer

##### Known PICI Mechanisms

PICIs (and phage satellites in general) are classic examples of phage-mediated horizontal gene transfer. After integration into the host genome via a PICI-encoded integrase, these islands lie quiescent until a lytic or temperate phage infects the cell pmc.ncbi.nlm.nih.gov. The phage’s replication cycle activates the PICI, leading to PICI excision and autonomous replication. The PICI then interferes with phage assembly to ensure its own DNA is packaged into viral capsids. The molecular details of this interference vary among satellite families and are often used to define them pmc.ncbi.nlm.nih.gov. For example, some PICIs encode a capsid morphogenesis protein that redirects phage capsid assembly to a smaller size, so that the phage genome no longer fits but the PICI genome does pmc.ncbi.nlm.nih.gov. Other PICIs instead produce terminase-mimicking sequences or small terminase subunits that preferentially package PICI DNA into phage heads pmc.ncbi.nlm.nih.gov. In all cases, the hijacked phage particles released upon cell lysis contain predominantly PICI DNA, which can infect new bacterial cells. This process is a specialized form of transduction, with PICIs effectively “piggybacking” on phage infections to transfer between hosts pmc.ncbi.nlm.nih.gov.

##### Phage Host Range Constraints

Under normal circumstances, a given PICI’s spread is limited by the host range of its helper phage and the specificity of its integration system. The integrase usually targets a particular attachment site in the host genome (often a conserved sequence or tRNA gene), and helper phages tend to infect only a narrow range of related bacteria. Consequently, most PICIs are confined to a single species or genus. For instance, SaPIs are mostly restricted to staphylococci, relying on *Staphylococcus* phages and specific chromosomal att sites. Similarly, the well-studied phage P4 satellite of *E. coli* depends on phage P2 and seldom ventures beyond *E. coli* and its close relatives pmc.ncbi.nlm.nih.gov. Thus, while PICIs significantly contribute to horizontal gene transfer, their movement typically occurs among closely related strains. Finding *the same* PICI in divergent bacterial species is unusual, suggesting an atypical transfer mechanism is at play.

##### Phage-Mediated HGT Overview

Beyond PICIs, phages mediate gene transfer through generalized transduction (packaging random host DNA) and specialized transduction (erroneous excision of prophage carrying adjacent genes). PICIs represent a more targeted system, carrying specific cargo genes (often virulence or resistance factors) that are deliberately mobilized by phage machinery pmc.ncbi.nlm.nih.gov. Importantly, many PICIs act as phage parasites – they usurp phage components for packaging and often interfere with phage reproduction, reducing phage burst size or blocking phage genome packaging pmc.ncbi.nlm.nih.gov. This antagonistic relationship can impose selective pressure on phages and hosts, generally constraining how far a PICI can spread. In summary, classical PICIs rely on precise integration and phage-hijacking strategies for HGT, which usually limit them to their original host species or close relatives.

#### Unique Features of Capsid-Forming PICIs (cf-PICIs)

Recent studies have identified a novel family of PICIs with properties that set them apart from typical phage satellites. **Capsid-forming PICIs (cf-PICIs)** encode their own capsid structural proteins and thus form PICI-specific viral heads, a capability not seen in other satellites <u>Pasteur.hal.science</u> pmc.ncbi.nlm.nih.gov. In essence, a cf-PICI carries all the genes necessary to build a small icosahedral capsid and package its DNA, requiring only the tail components from a helper phage to produce viable transducing particles <u>pasteur.hal.science</u>. This is in contrast to standard PICIs, which do not encode capsid or tail proteins – instead, typical PICIs supply at most a capsid size-modification protein and a small terminase, relying on the helper phage for all structural parts of the virion pmc.ncbi.nlm.nih.gov. The cf-PICI strategy represents a shift from hijacking phage capsids to manufacturing its own. Key distinctive features of cf-PICIs that may enable a broader host range include:

- **Self-Encoded Capsid Proteins:** Cf-PICIs encode the major capsid protein and assembly factors needed to form their own capsid shells <u>pasteur.hal.science</u>. They also encode a portal protein and head maturation protease, as inferred from conserved genes (e.g. a protease serine and HNH endonuclease found in cf-PICIs) pmc.ncbi.nlm.nih.gov. This autonomy in head formation means the PICI is not constrained to phage-specific head proteins. Any co-infecting phage can trigger cf-PICI packaging, since the PICI brings its own head components.
- **Head–Tail Adapter Modules:** Unique genes in cf-PICIs code for a head–tail connector and adaptor that allow the PICI’s capsid to attach to phage tail structures pmc.ncbi.nlm.nih.gov. In other words, the cf-PICI capsid is built with a “docking” system compatible with phage tails. This modular connector can interface with the tail tube or baseplate of the helper phage, effectively turning phage tails into delivery vehicles for PICI particles. The presence of these adapter proteins is a distinctive signature of cf-PICIs, absent in typical PICIs pmc.ncbi.nlm.nih.gov.
- **Complete Packaging Machinery:** Unlike standard PICIs which usually encode only a small terminase subunit (TerS) to hijack phage packaging, cf-PICIs encode a large terminase (TerL) as well pmc.ncbi.nlm.nih.gov. This implies that cf-PICI particles are packaged by a PICI-encoded terminase complex (TerS/TerL) that recognizes PICI genome pac sites and loads DNA into PICI capsids. By carrying a full complement of packaging genes, cf-PICIs ensure efficient encapsidation of their genome independent of the helper phage’s terminase. Essentially, the PICI’s DNA packaging is self-directed rather than parasitizing the phage’s packaging machinery.
- **Minimal Cost to Helper Phage:** Strikingly, cf-PICIs have been observed to have negligible impact on phage reproduction pmc.ncbi.nlm.nih.gov. Because they assemble separate capsids, they do not significantly compete for the phage’s head proteins or cause mispackaging of phage DNA. The helper phage can continue to produce its own full-sized virions in parallel. Empirical evidence from a prototype cf-PICI in *E. coli* (EcCI_EDL933_) showed that phage productivity was essentially unchanged in the presence of the PICI pmc.ncbi.nlm.nih.gov. In fact, the relationship may be near commensal or mutualistic rather than parasitic <u>pasteur.hal.science</u>. The ability to propagate without hurting phage fitness means phages carry cf-PICIs without selective pressure to eliminate them, facilitating persistent and widespread dissemination.
- **Broad Host Distribution:** Cf-PICIs have been identified in a wide array of bacterial species across both Gram-positive and Gram-negative phyla <u>Pasteur.hal.science</u> pmc.ncbi.nlm.nih.gov. Phylogenomic analyses indicate that cf-PICIs have emerged at least three times independently (convergent evolution) in different bacterial lineages <u>pasteur.hal.science</u>, underlining the adaptive value of this strategy. Moreover, surveys of bacterial genomes show that cf-PICIs are present in significantly more host species than typical PICIs or other phage satellites pmc.ncbi.nlm.nih.gov. For example, cf-PICIs occur in *Enterobacteriaceae* (e.g. *E. coli, Citrobacter*), *Pasteurellaceae* (*Haemophilus*), *Pseudomonadaceae* (*Pseudomonas*), and multiple Gram-positive families (Lactobacillaceae, Enterococcaceae, Bacillaceae, etc.) pmc.ncbi.nlm.nih.gov pmc.ncbi.nlm.nih.gov. Some of these genera had not been known to harbor any PICI-like elements until the discovery of cf-PICIs pmc.ncbi.nlm.nih.gov. This broad distribution suggests that cf-PICIs are less constrained by host specificity, consistent with an enhanced ability to move between species.
- **Conserved Integration and Regulation Modules:** Despite their novel structural genes, cf-PICIs retain the typical PICI modules for integration, replication, and regulation pmc.ncbi.nlm.nih.gov. They have a PICI-like integrase (usually a tyrosine recombinase) and repressor circuitry akin to other PICIs, and in some cases the entire left end of a cf-PICI (integrase through primase) is essentially identical to a standard PICI pmc.ncbi.nlm.nih.gov. This implies that cf-PICIs likely integrate into host genomes in a manner similar to other PICIs – often at specific attB sites such as tRNA or ribosomal gene loci that are conserved across many bacteria. By targeting a common site present in multiple species, a cf-PICI’s integrase could facilitate its establishment in diverse new hosts. Retaining these conserved modules means cf-PICIs can remain latent and regulated like normal PICIs, ensuring they don’t impose a burden on the host except during the phage-induced transfer cycle.

In summary, cf-PICIs combine a phage-like structural gene set (for capsid formation and DNA packaging) with the typical life-cycle modules of PICIs. This hybrid feature set is hypothesized to grant them a much broader host range. They are essentially modular mini-phages: lacking only tails and lysis functions, which they borrow from whatever phage infects the cell pmc.ncbi.nlm.nih.gov pmc.ncbi.nlm.nih.gov. These unique attributes likely underlie the observation that an identical cf-PICI can be found in multiple different bacterial species, a rarity for mobile genetic elements that are usually more restricted.

#### Interactions with Phage Tails and Bacterial Defenses

##### Phage Tail Hijacking

The hallmark of cf-PICIs is that they only require phage tails to form infectious particles <u>pasteur.hal.science</u>. During a helper phage infection, once the cf-PICI has replicated and assembled its capsids, it relies on available phage tail structures to complete the particle. Phage tail fibers and baseplates determine which bacteria a phage can infect (host range). By outfitting its capsid with a compatible head–tail connector, a cf-PICI can effectively hitch a ride on any tail produced by the helper phage pmc.ncbi.nlm.nih.gov. The result is a chimeric virion: a PICI capsid (containing PICI DNA) attached to a phage tail. Because the tail dictates host recognition, the cf-PICI particle will infect the same spectrum of hosts as the phage. If the helper phage has a broad host range (infecting multiple species or genera), the PICI will automatically share that broad host range. This is a critical point: the spread of cf-PICIs across species likely exploits phages that serve as “bridges” between different bacteria. Indeed, broad-host-range or generalized transducing phages could carry a cf-PICI from one species to another in a single transduction event. Even phages with a narrower range might facilitate stepwise transfer – for instance, a PICI could move from species A to closely related species B via one phage, and from species B to species C via another phage that infects B and C, and so on, eventually appearing in distantly related hosts. The modular nature of the cf-PICI capsid-tail interaction essentially decouples the PICI’s mobility from any one phage lineage, allowing it to opportunistically use whichever phage is present in a given host to propagate further.

##### Circumventing Bacterial Defenses

Moving between species means encountering different bacterial defense systems (e.g. restriction-modification systems, CRISPR-Cas immunity, and abortive infection mechanisms). How might cf-PICIs overcome these barriers? Several factors could facilitate their transit: (1) Small Genome and Phage-Like Packaging: The genome size of PICIs is relatively small (often 10–20 kb), which might reduce the chances of containing sites targeted by restriction enzymes compared to larger phage genomes. Additionally, if the helper phage modifies its DNA (e.g. glycosylation or methylation of phage DNA), the PICI genome packaged in the same cell may acquire similar modifications, helping it evade restriction systems in the new host. (2) Stealth and Low Immunogenicity: Because cf-PICI particles resemble phage particles externally, initial entry into a new bacterium via phage tail injection is likely to proceed as a normal infection. Once inside, the PICI DNA may integrate quickly into the chromosome via the integrase, potentially escaping detection. CRISPR-Cas systems that target known phage sequences might not recognize the PICI if its sequence is sufficiently diverged from common phage DNA. In fact, many PICIs share few homologous genes with intact phages pmc.ncbi.nlm.nih.gov, which could make them “invisible” to CRISPR spacers geared toward phages. Some PICIs even carry their own anti-defense genes; for example, certain PICI families encode proteins that confer anti-phage immunity to the host pmc.ncbi.nlm.nih.gov. A cf-PICI could theoretically harbor anti-CRISPR proteins or other defense inhibitors, ensuring its survival during transfer. (3) Mutualism with Phage: As noted, cf-PICIs impose little cost on phages and can even benefit them in some cases. One recent study showed an *E. coli* PICI providing superinfection immunity against the helper phage (preventing secondary phage infections) and also encoding a repressor that silences a large pathogenicity island, potentially freeing up host resources for phage replication pmc.ncbi.nlm.nih.gov. Such functions can make the PICI-carrying host more favorable for phage propagation. In a new species, these benefits could help the PICI establish by concurrently aiding any co-infecting phage, thus sidestepping abortive infection triggers (where bacteria commit cellular “suicide” upon phage attack) because the PICI might suppress lethal host responses or temper phage virulence just enough to avoid killing the host too fast. In short, cf-PICIs have the potential to buffer both phage and host interactions, smoothing the cross-species jump.

##### Possible Recombination Pathways

While phage-mediated transduction is the primary route for PICI transfer, we should consider if cf-PICIs might leverage other pathways. The excised form of a PICI is a circular DNA intermediate, analogous to a plasmid pmc.ncbi.nlm.nih.gov. Although not self-transmissible, there is a remote possibility that if a bacterium undergoes cell fusion or takes up DNA from the environment (natural competence), a cf-PICI could transfer without a phage. However, such events are likely rare compared to the highly efficient tail-mediated transduction. Another intriguing scenario is phage-phage recombination: if a cf-PICI capsid occasionally mis-packaged phage DNA or vice versa, hybrid particles or recombination between phage and PICI genomes could occur during co-infection. This might create novel mosaics or even broaden the PICI’s integration host range by acquiring multiple integrase specificities. While speculative, it underscores how cf-PICIs by virtue of sharing genomic space with phages could tap into diverse genetic exchange processes. Overall, the interactions of cf-PICIs with phage tails and host defenses paint the picture of a highly adaptable transfer mechanism – one that minimizes barriers and maximizes opportunities to invade new bacterial species.

#### Evolutionary and Ecological Implications

The ability of cf-PICIs to move horizontally across species boundaries has profound implications for microbial evolution and ecology. Below, we discuss several key impacts:

- **Accelerated Dissemination of Virulence and Resistance Genes:** PICIs often carry accessory genes that can enhance bacterial fitness or pathogenicity, such as toxin genes, antibiotic resistance determinants, or immune evasion factors pmc.ncbi.nlm.nih.gov. If a cf-PICI harboring a toxin gene (for example, the gene for a superantigen or a toxin in SaPIs) can transfer from one species to another, it can instantaneously bestow a new pathogenic trait on the recipient. This could lead to the emergence of new virulent strains in species that previously lacked that toxin. Similarly, antibiotic resistance genes embedded in a PICI could spread to completely different pathogenic species. In essence, cf-PICIs create a network of gene flow that links bacterial species into a shared genetic pool, potentially speeding up the spread of multidrug resistance or novel virulence factors across genus or family lines. This has obvious public health ramifications, as it means resistance or virulence traits are not confined to clonal expansion within one species but can hop into other opportunistic pathogens via phage-mediated transfer.
- **Breakdown of Species Barriers:** Traditionally, certain mobile elements (like conjugative plasmids) are known to move between species, but phage-related elements are often more restricted. Cf-PICIs blur this boundary. The fact that they have been found in at least 136 different species spanning Gram-positive and Gram-negative bacteria pmc.ncbi.nlm.nih.gov demonstrates an exceptional host range. Ecologically, this suggests that bacterial communities (for instance, in the human gut, soil, or marine environments) might be genetically more connected than anticipated. A cf-PICI can act as a shuttle vector, linking the evolution of one species to another. For example, environmental bacteria could donate a metabolic gene via a PICI to a pathogen under the right phage conditions, aiding the pathogen’s survival in a new niche. This inter-species mobility challenges the way we define species-specific gene pools and urges a perspective that considers the microbial community as a whole dynamic gene exchange network.
- **Phage–Satellite Co-evolution:** The emergence of cf-PICIs likely reflects a co-evolutionary balance between phages and satellites. In typical parasitic PICIs, the phage suffers a fitness cost, which can drive phages to evolve mechanisms to suppress or avoid PICIs, and PICIs in turn evolve counter-strategies. Cf-PICIs, by being nearly neutral or even beneficial to phages <u>pasteur.hal.science</u>, have changed this dynamic to a more cooperative one. The phage effectively becomes a willing carrier for the PICI. Evolutionarily, this could increase the longevity and reach of both entities: phages disseminating cf-PICIs might have an advantage if the PICI helps them (for instance, by providing superinfection immunity to prevent other phages from interfering pmc.ncbi.nlm.nih.gov). Over time we might see phage lineages that have adapted to always cart around a cf-PICI, essentially forming a phage-satellite symbiosis. Such phages would indirectly spread beneficial genes among hosts as a side effect of their life cycle. This co-driver relationship (as opposed to hijacker– hostage) is a fascinating shift, possibly explaining why cf-PICI lineages have been maintained over long evolutionary times and found in distant bacterial lineages (convergent evolution indicates this strategy is repeatedly favored <u>pasteur.hal.science</u>).
- **Impact on Bacterial Population Dynamics:** From a ecological standpoint, cf-PICI transfer could influence competition and cooperation in microbial communities. Bacteria acquiring a cf-PICI may gain competitive advantages, either through new traits or through the phage immunity that some PICIs confer pmc.ncbi.nlm.nih.gov. If a PICI provides immunity to its helper phage, it could create clonal lineages in a mixed-species community that are protected from certain phages, thereby shaping community composition. On the other hand, since cf-PICIs enable phage to infect more species without losing productivity, phages might broaden their host range in the presence of these elements. This could increase phage predation on non-host species once the PICI bridges them, linking the fate of different species under phage attack. In a clinical context, if one pathogen in a polymicrobial infection carries a cf-PICI, a therapeutic phage might inadvertently spread that PICI to co-infecting bacteria, disseminating genes of concern. Thus, understanding cf-PICI dynamics is important for phage therapy and microbiome intervention strategies.
- **Evidence of Recent Horizontal Transfer:** The discovery of near-identical cf-PICI sequences in different species is itself evidence of recent horizontal transfer in evolutionary terms. This highlights how quickly genetic elements can move in nature given the right conditions. It also raises questions about surveillance: these elements might be moving under our radar in various environments (hospital settings, agricultural sites, etc.). Monitoring for cf-PICI presence could become part of tracking the spread of resistance or virulence determinants. Their broad host range means that an outbreak of a resistant organism could potentially seed that resistance gene into completely different bacteria via a phage-mediated PICI transfer, complicating containment efforts.

In summary, the cross-species mobility of cf-PICIs represents a powerful mechanism for **genetic transduction across species boundaries**. It exemplifies nature’s innovation in gene transfer, merging phage infectious routes with plasmid-like specificity. Evolutionarily, it offers a route for genes to move not just vertically or by limited horizontal steps, but in quantum leaps across the tree of life of bacteria. Ecologically, it means communities are interlinked by these genetic exchange events, influencing traits like pathogenicity and antibiotic resistance in a web of interactions. The rise of cf-PICIs could be seen as nature’s response to selective pressures that favor genetic agility – whether in facing antibiotics, immune defenses, or nutrient challenges – by creating an element that can rapidly redistribute advantageous genes to where they are needed.

#### Conclusion

The same capsid-forming PICI appearing in multiple bacterial species can be explained by a novel mechanism of **phage tail-mediated modular transfer**. By encoding their own capsids and packaging systems, cf-PICIs liberate themselves from dependence on any single phage’s structural proteins and thereby gain access to a wider range of helper phages. This, combined with strategic use of conserved integration sites and minimal fitness costs to phages, enables cf-PICIs to travel across species via sequential phage infections. The proposed mechanism is supported by the unique genetic features of cf-PICIs (head-tail adaptors, terminase, etc.) and observations of their broad host distribution in nature pmc.ncbi.nlm.nih.gov. This cross-species horizontal transfer has likely contributed to the spread of important genes, linking bacterial populations that were once genetically isolated. Going forward, studying cf-PICIs offers insight into an evolutionary strategy that blurs the line between phage and mobile island, potentially informing new ways to control the spread of antibiotic resistance or to harness phages without unwittingly disseminating harmful islands. Understanding this mechanism in detail will be crucial for both fundamental microbiology and applied fields like infectious disease management, as it underlies a potent route of gene flow in the microbial world.

## Notes

### Competing Interest Statement

The authors have declared no competing interest.

